# Resolving the design principles that control postnatal vascular growth and scaling

**DOI:** 10.1101/2024.12.10.627758

**Authors:** Danielle Pi, Jonas Braun, Sayantan Dutta, Debabrata Patra, Pauline Bougaran, Ana Mompeón, Feiyang Ma, Stuart R Stock, Sharon Choi, Lourdes García-Ortega, Muhammad Yogi Pratama, Diomarys Pichardo, Bhama Ramkhelawon, Rui Benedito, Victoria L Bautch, David M Ornitz, Yogesh Goyal, M. Luisa Iruela-Arispe

**Author notes:** Lewis-Sigler Institute for Integrative Genomics, Princeton University, Princeton NJ, USA. These authors contributed equally (Danielle Pi, Jonas Braun).

## Abstract

After birth, tissues grow continuously until reaching adult size, with each organ exhibiting unique cellular dynamics, growth patterns, and (stem or non-stem) cell sources. Using a suite of experimental and computational multiscale approaches, we found that aortic expansion is guided by specific biological principles and scales with the vertebral column rather than animal body weight. Expansion proceeds via two distinct waves of arterial cell proliferation along blood flow that are spatially stochastic, yet temporally coordinated. Each wave exhibits unique cell cycle kinetics and properties, with the first wave exhibiting cell cycle durations as fast as 6 hours. Single-cell RNA sequencing showed changes in fatty acid metabolism concomitant with an increase in cell size. Mathematical modeling and experiments indicated endothelial cell extrusion is essential for homeostatic aortic growth and balancing excess proliferation. In a genetic model of achondroplasia, the aorta achieves proper scaling through enhanced cell extrusion while maintaining normal proliferation dynamics. Collectively, these results provide a blueprint of the principles that orchestrate aortic growth which depends entirely on differentiated cell proliferation rather than resident stem cells.

## INTRODUCTION

The design principles that coordinate the size of an adult organ and its scaling in relation to other tissues remain complex and elusive ^1^. At the cellular level, the basic determinants include spatiotemporally regulated increases in cell number and cell size. Nonetheless, the control knobs that mediate postnatal growth appear unique to a given organ, each with distinct cellular kinetics, proliferative patterns, and contributions from stem or non-stem cell sources ^2–4^.

In the vascular system, our understanding of collective growth properties of interconnected vessels first arose from studies in the endothelial lining of the retinal vasculature ^5^. In the retina, vascularization occurs radially from the central nerve. Expansion at the distal front occurs by angiogenesis, with tip cells advancing toward the periphery. Concurrently, the interconnected and more differentiated central vascular plexus shows active cell division and remodeling ^6^. This tissue’s stereotypical features, along with genetic tracing and proliferation analyses, revealed unequal replicative capacities across the continuum of vessel types (arterioles, capillaries, and venules). Specifically, endothelial cells from veins proliferate the most, followed by those located in capillaries that also overshadow the low proliferative capacity of endothelial cells from arterioles ^7^. Despite clear inequalities in division kinetics, active cellular rearrangements and migration ultimately yield a balanced and perfectly scaled vascular network. This occurs because cells originating in veins actively migrate against blood flow through interconnected tubes across capillaries to establish residency in arteries ^7^. These and other studies implied that arterial cells exhibit a greatly reduced ability to proliferate and that the expansion of arteries to scale the vascular network is accomplished by the proliferation of endothelial cells from veins ^8^. In adult zebrafish, for example, the repair of microvascular networks following fin amputation occurs through the local proliferation of endothelial cells in veins, which then reconstitute both capillaries and arteries ^7^. Collectively, these studies suggested that quiescence is a requirement for the acquisition of arterial fate and further, that the arterial cell fate is incompatible with proliferation ^9,10^. The notion that arterial cells cannot proliferate establishes a difficult paradox: How do large arteries, far removed from venules, grow postnatally in length and width?

In many ways, the aorta does not fit the mold of most arteries studied thus far. The dorsal aorta is the largest artery in the body and is farthest removed from capillaries and veins, leaving cellular migration from other vascular beds an unlikely source of new cells. Furthermore, unlike other vessels, the aorta emerges by the direct differentiation of endothelial cells from mesoderm in a process referred to as vasculogenesis ^11^. Importantly, the aorta and its branches are populated by fully differentiated arterial endothelial cells, having acquired arterial fate even prior to birth ^12^. In addition, this vessel is exposed to the highest levels of shear stress and physical forces associated with hemodynamic flow ^13^. The process by which the aorta gains additional cells and scales to the growing body to comply with the metabolic needs of peripheral organs remains unresolved.

Adopting a highly interdisciplinary and multiscale approach, we uncovered that postnatal growth of the aorta occurs monotonically through coordinated *in situ* proliferation and enlargement of fully differentiated arterial endothelial cells. We found that major and minor cellular axes generally follow the aortic axial and circumferential growth dynamics. The integrated parameters of proliferation, cell size, and cell division orientation fully account for tissue expansion in length and diameter. Interestingly, postnatal proliferation of the aortic endothelium occurs in two distinct temporal waves. Quantitative clone tracing within and across each wave revealed distinct and highly reproducible features of the waves across animals and sexes. The first wave exhibits fast cell cycle kinetics (as short as 6 hours) and asynchronous division, while the second wave is slower and occurs synchronously. Importantly, our mathematical model of aortic growth revealed that cell proliferation alone significantly overshoots the number of required cells. The model predictions led us to experimentally uncover a surprising mechanism to recover cell number balance, which requires a high level of physiological endothelial cell extrusion into the circulation. This mechanism of cell extrusion enables the aorta to scale appropriately to the size of the vertebral column, as revealed by data from a disease model of achondroplasia. In this matter, the principles that underlie proliferation are fixed, while cell extrusion is dynamic and adaptable. Our observations, otherwise not possible without the quantitative multidisciplinary approaches applied here, uncover several unexpected paradigms underpinning tissue growth and organ scaling, with broad implications for the field of vascular biology and beyond.

## RESULTS

### Longitudinal and circumferential growth dynamics are uncoupled

Our initial goal was to create a quantitative framework to study the postnatal growth of the aorta. Using microcomputed tomography (microCT), we measured the dimensions of aorta *in situ* from birth to adulthood (**Figure 1A, Video S1**). Our analysis focused on a well-defined thoracic segment of the aorta comprising the region between the first and eighth pair of intercostal arteries (**Figure 1B**). This region is relatively straight, allowing us to measure relevant geometric features in animals over time and space. Using this framework, we investigated whether the aorta’s length and circumference—its two key features—grow in parallel or follow distinct trajectories. We found evidence for the latter. The aorta’s length grew in two phases: minimal growth from postnatal day 0 to 7 (P0 to P7), followed by substantial growth from P7 to P30 (**Figure 1C, S1A**). In contrast, the circumference grew steadily from P0 until reaching maturity around P21 (**Figure 1D**). We then examined whether the aorta’s length or circumference scaled with animal weight. Surprisingly, we found no clear correlation (**Figure 1E,G**), and very different kinetics between length/circumference and weight when considered on an absolute scale (**Figure 1G** inset); however, aortic volume scaled at a more similar fold-change to animal weight (**Figure S1C**). These results suggest that aortic growth dynamics reflect a more regulated process rather than simply scaling with overall animal weight.

**Figure 1.**
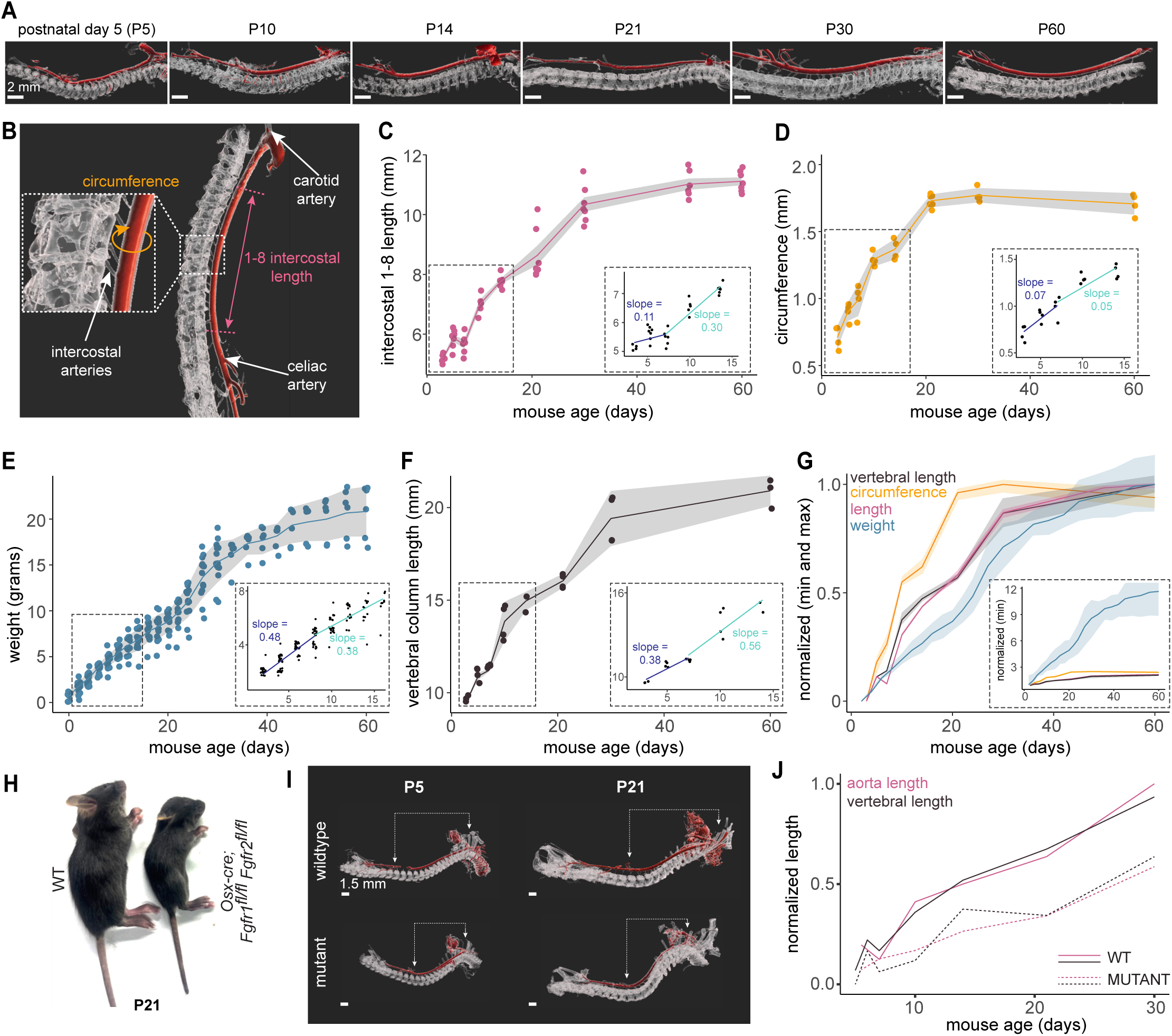
Aortic length scales with the vertebral column. (A) Microcomputed tomography reconstructions showing the aorta (red) and vertebral column (white). Scale bar = 2 mm. (B) Annotated anatomical landmarks of the descending aorta. (C) Measured length (mm) of descending aorta between the 1st - 8th intercostal vessels. Inset shows the rates of growth of aorta length from postnatal day 3 to 7 (P3-P7) and from P7 to P14. Each dot represents a biological replicate, with the mean and standard deviation shown. N = 5-7 animals per age. (D) Measured circumference (mm) of the descending aorta. Inset shows the rates of growth of circumference before and after P7. Each dot represents a biological replicate, with the mean and standard deviation shown. N = 3-6 animals per age. (E) Measured weight of animals (g). Inset shows the rates of growth of weight before and after P7. Each dot represents a biological replicate, with the mean and standard deviation shown. N = 6-28 animals per age. (F) Measured vertebral column length (mm) between T1 and L1. Inset shows the rates of growth of vertebral column length before and P7. Each dot represents a biological replicate, with the mean and standard deviation shown. N= 2-4 animals per age. (G) Compilation of vertebral column length (black), aorta circumference (orange), aorta length (pink), and animal weight (blue), normalized to minimum and maximum values of averages for each condition. Inset shows measurements normalized to minimum values. The mean and standard deviation is shown. (H) Representative images of wildtype (WT) and mutant (*Osx-cre; Fgfr1^fl/fl^ Fgfr2^fl/fl^*) animals at P21. (I) Representative microcomputed tomography reconstructions of wildtype and mutant (*Osx-cre; Fgfr1^fl/fl^ Fgfr2^fl/fl^*) aorta and vertebral column. The arrows point to the subclavian artery branch point at the arch, and the celiac trunk branch point, respectively. Scale bar = 1.5 mm. (J) Aorta (pink) and vertebral (black) length of wildtype and mutant (*Osx-cre; Fgfr1^fl/fl^ Fgfr2^fl/fl^*) animals, normalized to minimum and maximum values of averages for each condition. N = 3-5 animals per age, per condition.

Given the close anatomical relationship between the aorta and the vertebral column, we investigated whether the length of thoracic aorta corresponded with the length of the thoracic vertebrae, or if the aorta grew independently as, in humans, intercostal pairs stereotypically branch off at corresponding vertebral levels ^14^. Thus, we used microCT to determine the correspondence between vertebral levels and aortic branching landmarks in mice. We found that across all ages and biological replicates, the two aorta landmarks of the subclavian artery and celiac trunk branch points were stereotypically associated with the same vertebral levels (T2-T3 and L1-L2, respectively) (**Table S1**). Notably, the growth of vertebral length scaled closely with aorta length (**Figure 1F,G, S1B**). These results demonstrate that unique growth kinetics underlie aorta development, with longitudinal growth being highly coordinated with vertebral growth.

### Aortic length scales with the vertebral column

Given the strong concordance between the aorta and vertebral length, we questioned whether the scaling relationship is conserved in scenarios where the spinal growth dynamics are affected. To address this question, we used a mouse model of achondroplasia (*Osx-cre; Fgfr1^fl/fl^ Fgfr2^fl/fl^*). In this model, chondrogenesis and bone growth are affected through genetic deletion of *Fgfr1* and *Fgfr2* in the osteoprogenitor lineage ^15^, providing an excellent platform to probe how the aortic length is affected (**Figure 1H**). While diminished spine growth in this model has previously been reported ^15^ (**Figure S1D**), our imaging results from microCT indicated a correspondingly shorter aorta compared to wildtype throughout postnatal growth (**Figure 1I**). Importantly, besides the shorter length, the aorta growth kinetics also scaled closely with the stunted spinal growth (**Figure 1I,J**). These results reinforced a precise and temporal scaling relationship between the thoracic aorta and vertebral column that is preserved even under disease settings.

### Endothelial cell size increases with age

To deconvolve the relationship between cellular- and tissue-level growth, we focused on the inner cellular layer of the aorta (the endothelium) as the central experimental paradigm. For this, we excised the aorta for *en face* whole-mount imaging of the endothelium and downstream analysis of cell number, size, density, and shape over time (**Figure 2A, S2A**). Conventional intensity-based computational methods of cellular segmentation performed poorly on the aorta images due to the thin endothelial layer and unavoidable visualization of underlying smooth muscle cells. Thus, we developed a custom deep-learning image analysis pipeline with numerous quality control steps to automatically and reliably segment nuclear and cellular features of the endothelium at scale (**Figure 2B, S2C; STAR Methods**). We found that the length of the aorta between intercostals 1-8 increased 2.4 fold, from an average of 5.19 mm in P3 to average of 12.31 mm in P60, whereas endothelial cell number in this region increased by 2.8 fold, from an average of 18,411 to 50,827 (**Figure 2C**). We next asked whether this increase in cell number was coupled with tissue expansion, such that the cell density remains constant, or whether one aspect superseded the other. The measurements revealed cell density along the length and circumference to peak in the first seven days and then decrease over time. Specifically, cell density went from an average of 4,440 cells/mm^2^ at P3-7 to an average of 2,320 cells/mm^2^ after P30. This implied an unequal relationship where increase in cells superseded the growth in surface area early on, resulting in a higher cell density during the early time points (**Figure 2D,E**). We next asked whether the increase in cell number occurred in an isotropic manner or at specific spatial foci, and found proliferation to be uniform throughout the aorta (**Figure 2D,E**). As density decreased P7 onward but aortic growth continues to occur, we reasoned that cell size must increase to account for reduced density over time. Using Pecam labeling to mark endothelial cell borders, we indeed found that cell area increased significantly from P7 onwards (p value = 0.0055, t-test, two-sided) (**Figure 2F**). Cell height did not change appreciably during this time window (**Figure S2B**). However, as was the case for global aorta measurements, the dynamics of cell length (major axis of an approximated ellipsoid) was distinct from the cell width (minor axis) (**Figure 2G-I**). With these rich cell-level measurements, we asked whether we could recapitulate the global longitudinal and circumferential dynamics of the whole tissue. We computed the average number of cells along the length (or circumference) axis (**STAR Methods; Figure S2D-F**), and multiplied this with the length of major (or minor) axis to estimate the aorta length (or circumference) and found the estimations to closely match the measured values (**Figure 2J-L**).

**Figure 2.**
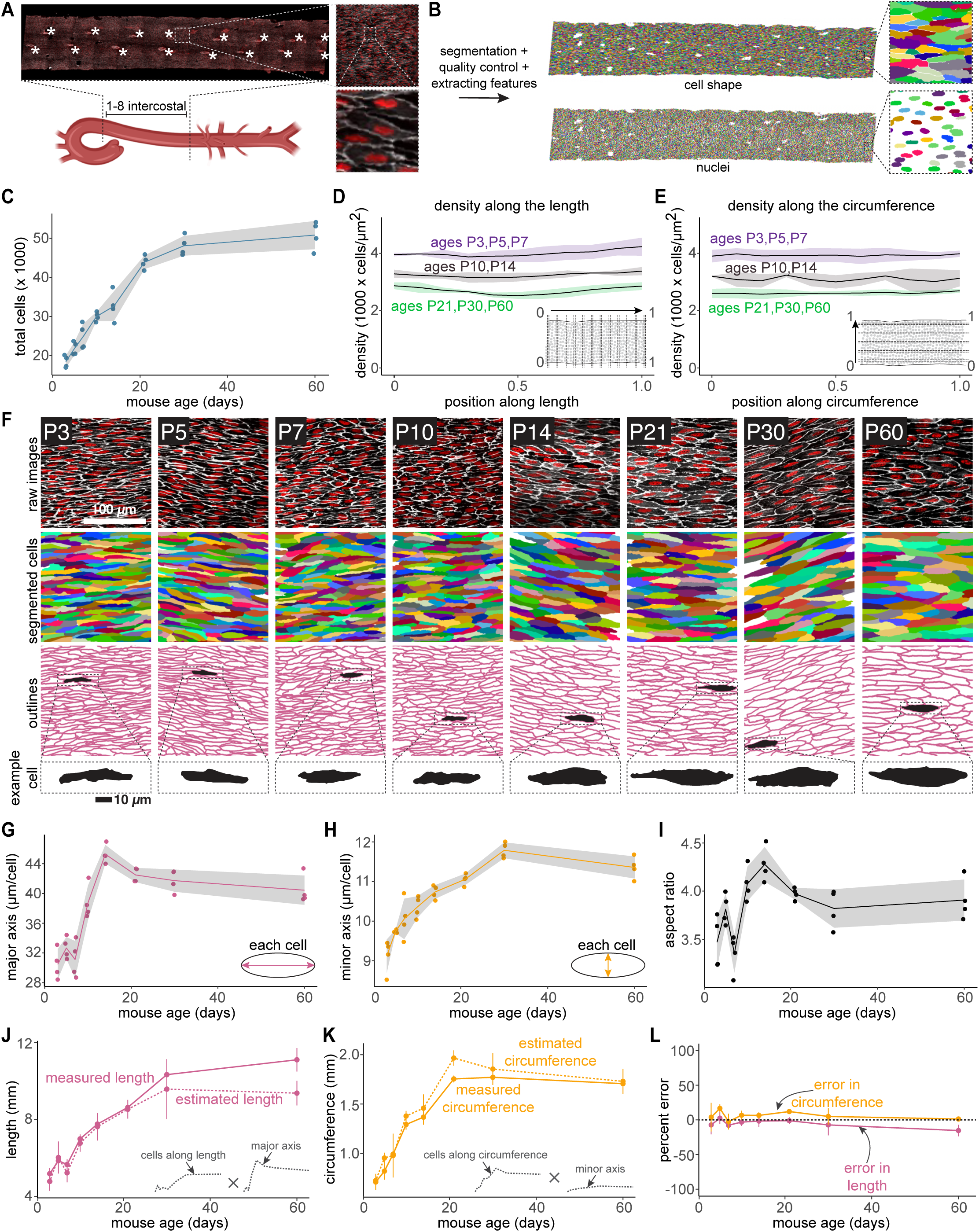
Dynamics of endothelial cell size during postnatal development. (A) *En face* whole mount image of an aorta between the 1st and 8th intercostal, immunostained for ERG (nuclei, red) and PECAM (borders, white). Asterisks indicate intercostal artery branches within the aorta. (B) Work flow of whole mount image processing. Machine learning segmentation was used to extract cell shape and nuclei of all endothelial cells in the thoracic aorta. Each segmented cell and nuclei was assigned a randomized color. (C) Total number of endothelial cells between the 1st - 8th intercostal vessel in the descending aorta. Each dot represents a biological replicate, with the mean and standard deviation shown. N = 4 animals per age. (D) The density of cells (1000 cells per µm^2^) along the length of the thoracic aorta. Position 0 refers to the cranial direction and position 1.0 refers to the caudal direction. The mean is denoted by the solid line and shading indicates standard deviation. N = 4 animals per age. (E) The density of cells (1000 cells per µm^2^) along the circumference of the thoracic aorta. Position 0 refers to the left of an en face aorta and position 1.0 refers to the right. The mean is denoted by the solid line and shading indicates standard deviation. N = 4 animals per age. (F) Representative images of *en face* whole mount immuno-stained endothelium within the aorta (red: ERG; white: PECAM) and machine learning segmentation of individual cells, where each segmented cell has a randomized color. A representative cell from each age is shown in the last row in black. (G) Average major axis (µm) of all cells segmented from each aorta. Each dot represents a biological replicate, with the mean and standard deviation shown. N= 283 - 34434 cells per aorta, 4 animals per age. (H) Average minor axis (µm) of all cells segmented from each aorta. Each dot represents a biological replicate, with the mean and standard deviation shown. N = 283 - 34434 cells per aorta, 4 animals per age. (I) Average aspect ratio of all cells segmented from each aorta. Each dot represents a biological replicate, with the mean and standard deviation shown. N= 283 - 34434 cells per aorta, 4 animals per age. (J) The estimated length of the aorta (dashed line) is calculated by multiplying the average number of cells along the length by the average major axis. The experimentally measured length of the aorta (solid line) is shown for comparison. Each dot represents the mean length, with error bars showing the standard deviation between biological replicates. N = 4 animals per age. (K) The estimated circumference of the aorta (dashed line) is calculated by multiplying the average number of cells along the circumference by the average minor axis. The experimentally measured circumference of the aorta (solid line) is shown for comparison. Each dot represents the mean length, with error bars showing the standard deviation between biological replicates. N = 4 animals per age. (L) Percentage error between measured and estimated length (pink line), and percentage error between measured and estimated circumference (orange line). Each dot represents the mean percentage error, with error bars showing the standard deviation between biological replicates.

### Aortic growth is accomplished by collective and egalitarian proliferation of differentiated cells that occurs in two waves

Given the monotonic increase in total cell number, we assessed whether proliferation occurred in specific locations, and if this was steady or in bursts by measuring the fraction and distribution of cells positive for Ki67, a proliferation marker (**Figure 3A, S3A, Table S2**). Surprisingly, we found that endothelial cell proliferation occurred in two waves, with the first wave lasting until P10, followed by a short quiescent phase before another proliferative wave peaked around P21 (**Figure 3B, S3B**). The proliferation was negligible by P60 (ranging from 1.5 to 2.2%), consistent with the saturation of cell number (**Figure 2C**). In line with cell density calculations, where we found an isotropic increase, the Ki67-positive cells were uniformly distributed throughout the aorta (**Figure 3C,D**). Given the unexpected observation of biphasic Ki67-positive dynamics, we confirmed this finding by quantifying the number of cells with active mitotic spindle (acetylated α-tubulin) as another proxy for cells undergoing division (**Figure S3D**). Here again, two peaks of total cells with active mitotic spindles were found, each mirroring those seen for Ki67-positive dynamics (**Figure 3E**). Next, we asked whether the cells undergoing division do so relative to a particular direction of blood flow or if the direction is decided stochastically. By measuring mitotic spindle angles (**Figure 3F, S3C**), we found cell division to occur predominantly along the direction of blood flow (0°). Cell division orientation therefore largely contributed to addition of cells in the longitudinal direction, consistent with previous work performed in small vessels ^16^. Importantly, the range and distribution of angles, from +30 to -30 degrees relative to the direction of flow, contributed to increase in the circumference (**Figure 3G**). Intriguingly, the proliferative waves and the growth in aorta length did not always occur in overlapping time windows, suggesting regulation between balancing growth versus proliferation.

**Figure 3.**
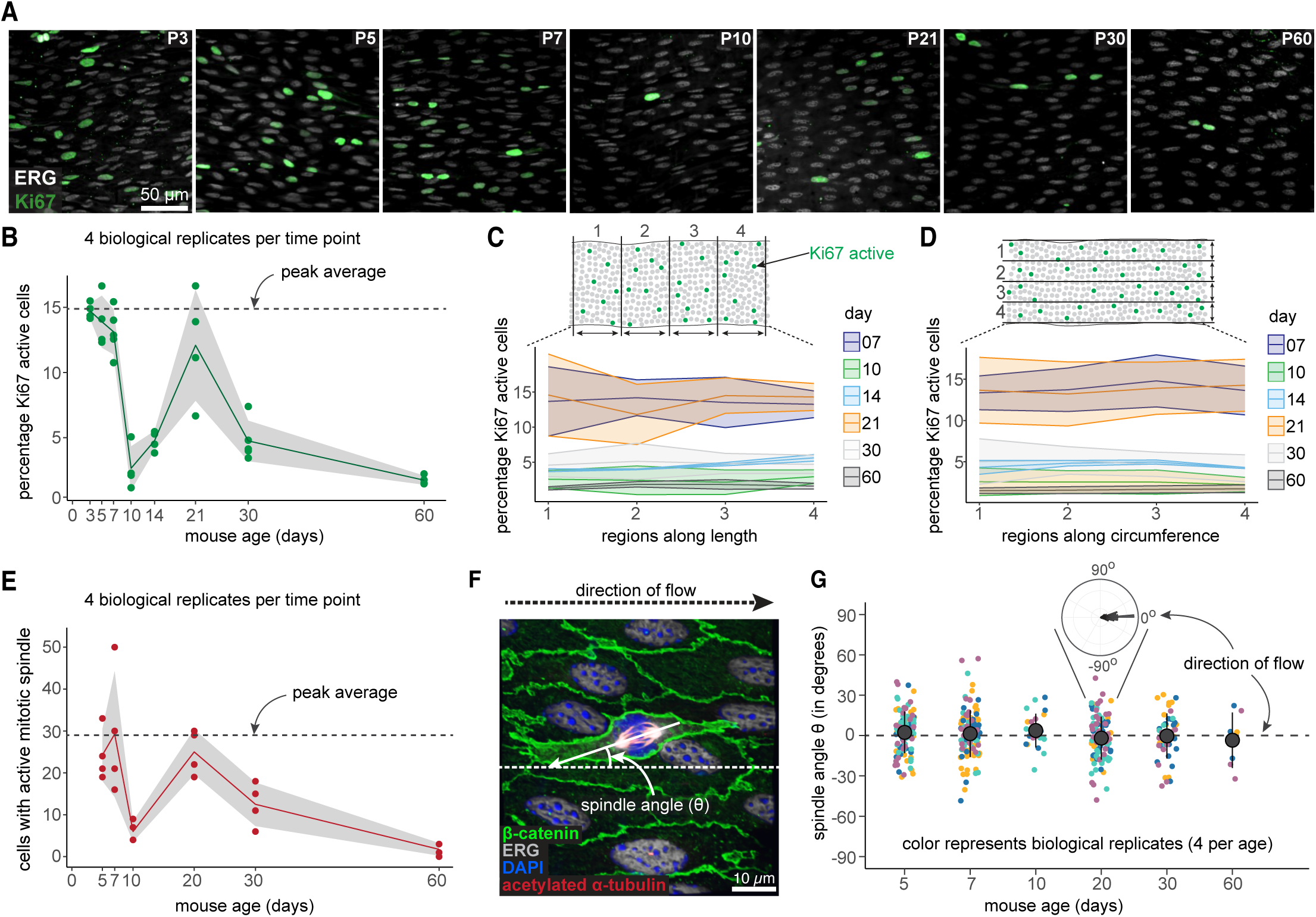
Endothelial cell proliferation occurs in two waves during aortic growth. (A) Representative images of *en face* whole mount immuno-stained aortas showing proliferation in the endothelium (white: ERG; green: Ki67). Scale bar = 50 µm. (B) Percentage of Ki67 active cells from all segmented endothelial cells from each aorta. Each dot represents a biological replicate, with the mean and standard deviation shown. N = 3980 - 69827 cells per aorta, 4 animals per age. (C) Percentage of Ki67 active endothelial cells within regions along the length of the aorta. Position 1 refers to the cranial direction and position 4 refers to the caudal direction. The mean is denoted by the solid line and shading indicates standard deviation. N = 19616 - 69827 cells per aorta, 4 animals per age. (D) Percentage of Ki67 active endothelial cells within regions across the circumference of the aorta. Position 1 refers to the left of an *en face* aorta and position 4 refers to the right. The mean is denoted by the solid line and shading indicates standard deviation. N = 19616 - 69827 cells per aorta, 4 animals per age. (E) Count of endothelial cells with active mitotic spindles (determined through acetylated α- tubulin immuno-staining) from each aorta. Each dot represents a biological replicate, with the mean and standard deviation shown. N = 4 animals per age. (F) Representative image showing whole mount immuno-staining of an active mitotic endothelial cell (green: β-catenin; white: ERG; blue: DAPI; red: acetylated α-tubulin). The spindle angle (θ) is the angle between the mitotic spindle and direction of flow. Scale bar = 10 µm. (G) Distribution of endothelial mitotic spindle angle (θ) from each age relative to the direction of flow. Each dot represents a cell with an active mitotic spindle and each color represents a biological replicate. Error bars are mean +/- standard deviation. N = 4 animals per age.

To orthogonally support or refute the two waves of proliferation and to ascertain the cell identity of the proliferating cells, we leveraged single-cell RNA sequencing (scRNA-seq). Four sequencing libraries were generated from aortae at P5, P10, P20, and P30, and quality controlled for downstream analysis (**Figure 4A**, **Figure S4A-D**). Using classical markers, we identified five major cell clusters: endothelial, smooth muscle, fibroblasts, inflammatory cells, and proliferating cells (**Figure 4B,C**). Due to the aorta’s small size at this age, endothelial cells from P5 and P10 aortae were isolated by whole-tissue digestion. In contrast, cells from P20 and P30 aortae were isolated by our standard surface trypsinization method ^17^. The difference in cell isolation method explains the abundance of fibroblasts and vascular smooth muscle cells at the early time point libraries and the greater enrichment of endothelial cells in the P20 and P30 libraries (**Figure 4A**). Despite these differences, after quality control, we were able to evaluate 17,516 endothelial cells (2,150 at P5, 1,273 at P10, 5,071 at P20 and 9,022 and P30). The combined cellular constituencies displayed the anticipated populations and their predicted abundances ^17,18^. Using *MKi67, Top2a,* and *Ube2c*, we found that all proliferating cells were segregated from the rest, forming a unique cell cluster that included endothelial, fibroblast, and smooth muscle cells (**Figure 4D**). Despite their proliferative identity, distinct cell types could still be identified based on their canonical markers (*Pecam1* and *Cdh5* in the case of the endothelium) (**Figure 4D, insert and C**).

**Figure 4.**
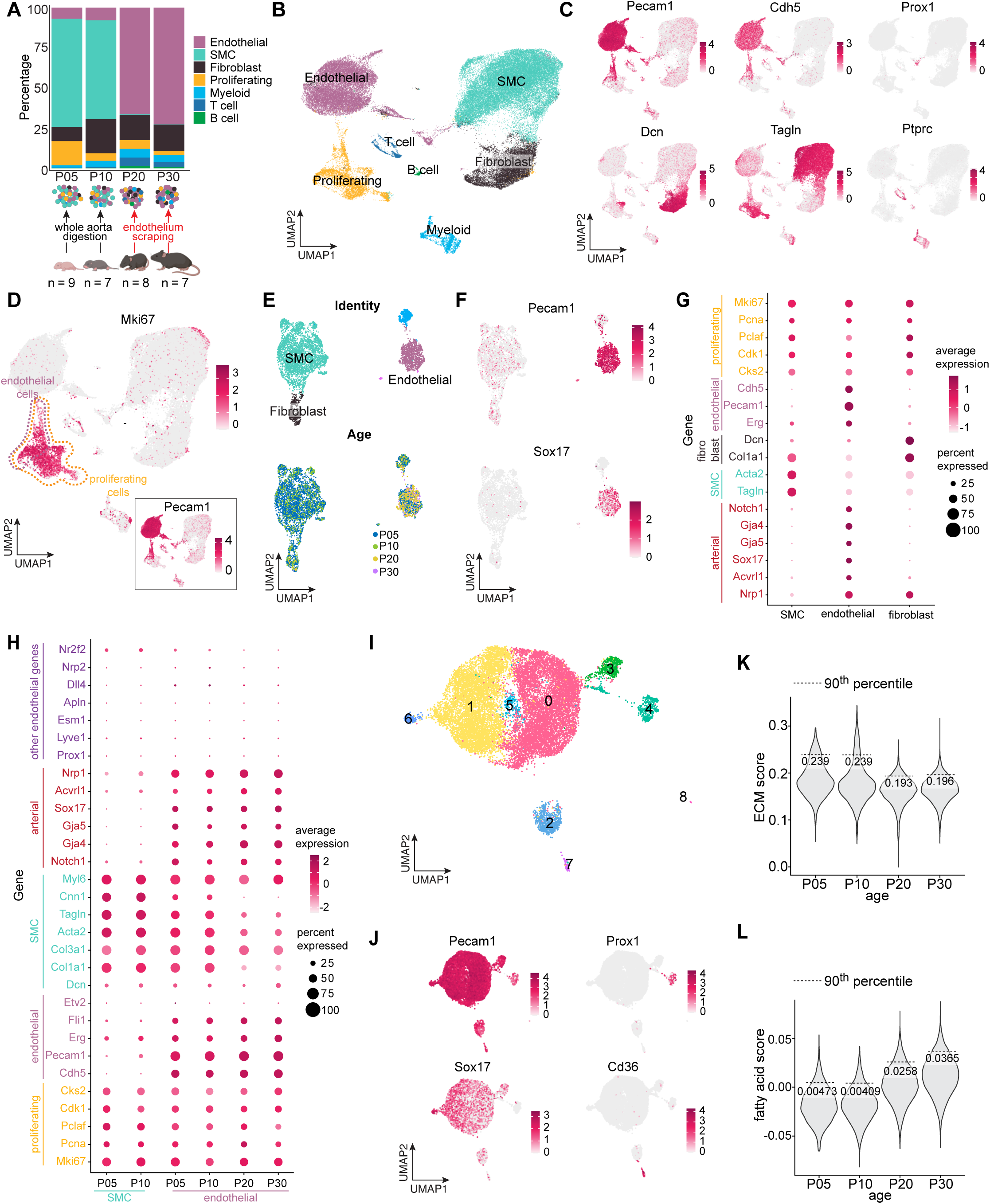
Proliferative endothelial population retains arterial fate. (A) Schema of the approach and number of cells obtained. (B) UMAP of combined libraries after quality control. (C) Cell identity markers for endothelial cells (*Cdh5*, *PECAM*, *Prox1*), fibroblast (*Dcn*), vascular smooth muscle cells (*Tgln*), and inflammatory cells (*Ptprc*). (D) UMAP of all cell populations identifying proliferative cluster (yellow outline). The endothelial proliferative population is outlined in purple and is based on the expression of *Pecam1* (inset). (E) Subsample of proliferating cells in UMAPs showing clusters (upper) and origin (bottom). (F) Cell identity markers demonstrating the endothelial cluster (as per *Cdh5* and *Sox17* expression). (G) Dot plot illustrating levels of expression of markers for: arterial, smooth muscle cell (SMC), fibroblast, pan-endothelial, and proliferating cells. The color of each dot indicates average expression and the size of each dot indicates percentage of cells that express each gene. (H) Dot plot of proliferating cells in P5 and P10 smooth muscle cells (SMC), and in P5, P10, P20, and P30 endothelial cells. The markers are identical to those in G. The color of each dot indicates average expression and the size of each dot indicates percentage of cells that express each gene. (I) UMAP of and cluster distribution for all endothelial cells (non-proliferating) from all ages. (J) UMAPs of major endothelial clusters identified by *Pecam1*, *Prox1*, *Sox17*, and *CD36* expression. (K) Violin plots of extracellular matrix (ECM) score from P5, P10, P20, and P30 endothelial cells. The score of the 90^th^ percentile is indicated in numbers and with a dashed line. (L) Violin plots of fatty acid score from P5, P10, P20, and P30 endothelial cells. The score of the 90^th^ percentile is indicated in numbers and with a dashed line.

Further evaluation of the proliferative population allowed us to gain a more refined resolution into the three cell groups, identified as fibroblasts, smooth muscle cells, and endothelial cells using standard markers (**Figure 4E,F**). Importantly, additional appraisal of the 1,031 proliferative endothelial cells revealed the unexpected retention of arterial identity, as per expression of *Notch1, Gja4, Gja5, Sox17, Acvrl1,* and *Nrp1* (**Figure 4G**). This prompted the question of whether arterial identity was equally preserved in proliferating cells across all time points, as cell cycle exit was previously reported as a prerequisite for arterial specification ^8,10^. Indeed, we found the same pattern of expression during all the time points, demonstrating retention of arterial identity while these cells were proliferating with equivalent levels of expression across all developmental time points (**Figure 4H**). We also evaluated canonical vein, capillary, and lymphatic markers, and observed no gain of expression in those transcripts that would otherwise indicate a change in cell fate in the proliferating population (**Figure 4H**). Using this dataset, we further determined the percentage of proliferating cells per time point, confirming, once again, the presence of two waves of proliferation (**Figure S4E**). Specifically, we found that 17.63% of endothelial cells were proliferating at P5 with a drastic reduction to 4.48% at P10, followed by a significant increase to 6.2% by P20, and a decline thereafter (3.1% by P30). The findings from the Ki67 immunofluorescence and the scRNAseq are concordant, supporting the presence of two proliferative waves. The difference between the absolute (not relative) percentage of proliferating cells by scRNAseq and the Ki67 immunofluorescence quantification is not surprising, as we had to use different dissociation and cell recovery process across the waves due to technical limitations. Moreover, Ki67 protein only captures cells activated and primed to divide, while scRNAseq is highly subsampled and does not capture all G1/Ki67 positive cells. Such a possibility would suggest a longer G1 in the second wave, a hypothesis we later pursue with experimental confirmation.

The transcriptomic data also allowed us to explore the progressive changes associated with non-proliferative endothelial cells from P5-P30. Thus, we subsampled these cells (**Figure 4I**) and identified a very large arterial endothelial population, in addition to lymphatic endothelial cells (as per Prox and Lyve1) and capillaries (as per CD36) from the adventitia (**Figure 4J**). Assessment of unique transcripts between all four age groups revealed distinct maturation stages despite retention of endothelial and arterial fate. In particular, we found a progressive increase of genes associated with ion channels, Akt activation, calcium dynamics, gain of histocompatibility genes, and changes in metabolism (**Figure S4J)**. Importantly, we also found a decrease in overall matrix production over time (**Figure 4K, S4I**), but a distinct and rather impressive increase in transcripts associated with fatty acid metabolism (**Figure 4L, S4H),** while there was no change in Acetyl CoA metabolism across the ages **(Figure S4F, G).**

To gain additional granularity on the clonal proliferation dynamics, we performed clonal analysis using the Rainbow system (mCerulean, mCherry, or mOrange). This approach reports on proliferation, but also adds information on the preferred direction of endothelial cell division as it’s translated into clonal outgrowth of cells along the length of the aorta. If individual migration were prevalent, one would expect the three reporter colors to be interspersed. Conversely, if no migration or only collective migration were present, cells sharing the same color would remain together. We validated these assumptions in three vascular beds—two large vessels (aorta and vena cava) as well as small vessels in the retina from the same animals. Within large vessels, clones were found to be spatially localized (**Figure 5B, S5A**). In contrast, retinal endothelial cells exhibited interspersion of different colors, suggesting individual cell migration consistent with the literature ^19^ (**Figure 5B**). Furthermore, aortic endothelial cell clones aligned primarily in the direction of flow, mimicking what was observed with cell-division orientation (**Figure S5A**), whereas the endothelial cell clones in the vena cava expanded in both the longitudinal and lateral directions. Importantly, these results established the feasibility of performing quantitative clonal analysis over time. The findings also highlighted the power of this quantitative platform in establishing the complex and rationalizable connections between global and local measurements, while revealing unexpected biphasic proliferation kinetics.

**Figure 5.**
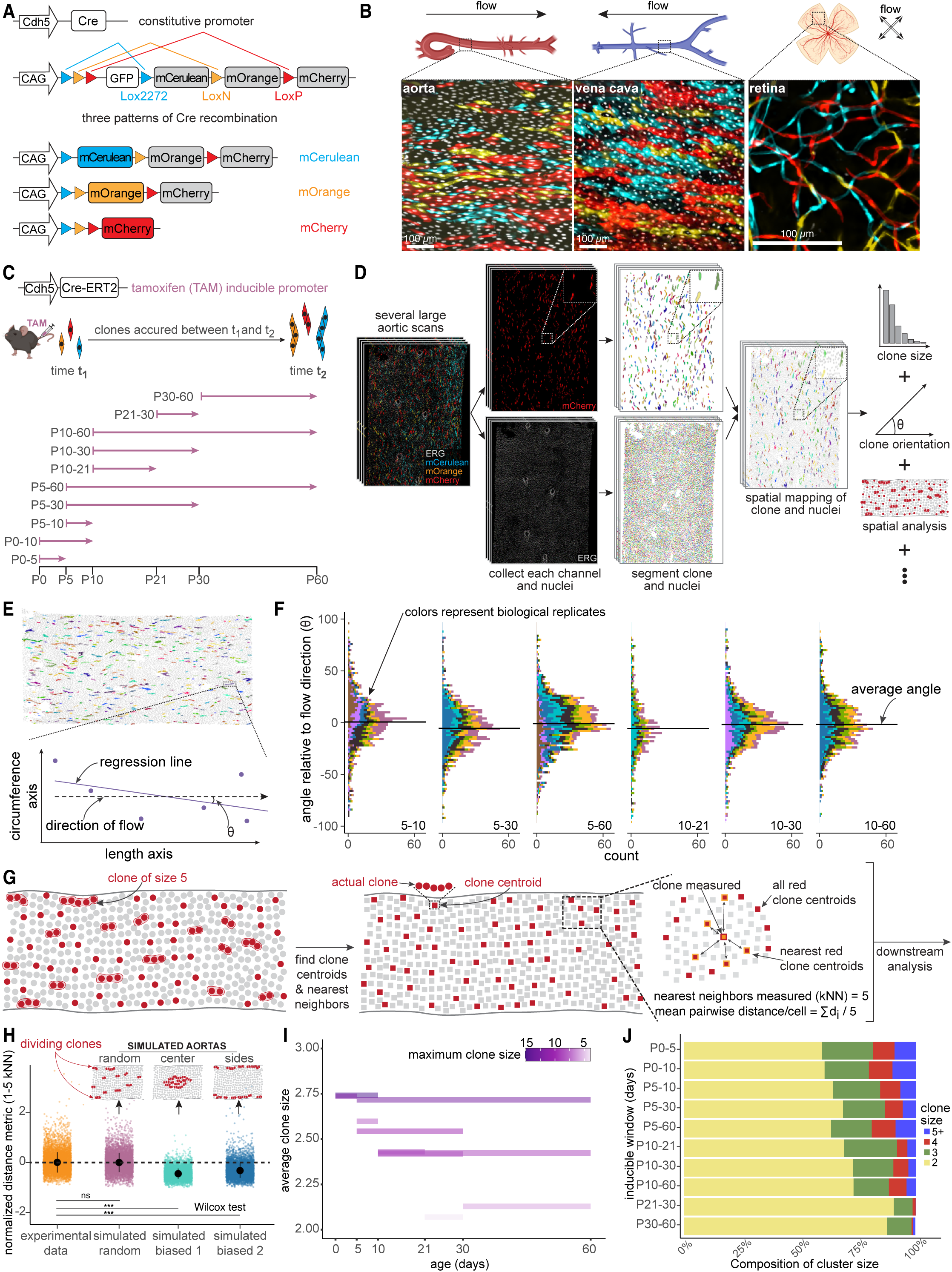
Endothelial clonal dynamics is spatially uniform but temporally regulated across the two proliferative waves. (A) Schematic of the constitutive rainbow lineage tracing animal. Cre expression is driven by the constitutive Cdh5 promoter, which leads to random recombination at Rainbow^fl/+^ as early as E7.5; recombined cells either express mCerulean, mOrange, or mCherry fluorescent protein. (B) Representative images showing descending aorta, infrarenal vena cava, and retina whole mount immuno-stains (white: ERG; blue: mCerulean; yellow: mOrange; red: mCherry) from constitutive *Cdh5-Cre Rainbow^fl/+^* animals. (C) Schematic of the tamoxifen (TAM) inducible lineage tracing animal, and times of induction and harvest. At time of induction, recombined cells either express mCerulean, mOrange, or mCherry fluorescent protein and cells accrue as clones until time of harvest. (D) Work flow of machine learning image segmentation of rainbow clones. (E) Method of calculating clone orientation. A regression line is fitted through the centroid points of all members of a clone, and the angle between the regression line and the direction of flow (θ) is taken as the angle of clone orientation. (F) Clone orientation from postnatal day 5-10, 5-30, 5-60, 10-21, 10-30, and 10-60. The overall angle (θ) of each clone relative to the flow direction is shown, with angle on the y-axis (in degrees) and frequency on the x-axis. Each bar represents a clone and each color represents a biological replicate. The average angles across all clones from all biological replicates are indicated by the black line. N = 5-8 animals per time point. (G) Schematic of clone spatial analysis. The centroid is identified for each clone, then nearest neighbor analysis is performed between the centroids of clones. (H) 1000 simulations are performed for random (simulated random), center biased (simulated biased 1), or side biased clone distribution (simulated biased 2), then compared to real data for nearest neighbor analysis. Error bars are mean +/- standard deviation. (I) Average clone size of non-singlets from each time window; the color of the bar represents the maximum clone size found within that time window. N = 5-8 animals per time point. (J) Proportion of cluster sizes of non-singlets, found across all induction and harvest time windows.

### Endothelial clonal dynamics is spatially uniform but temporally regulated across the two proliferative waves

Establishing endothelial clones as cells underwent division enabled subsequent systematic, quantitative, and comparative probing of clonal dynamics by using an inducible model to resolve features across time. Using tamoxifen-inducible *Cdh5 Cre-ERT2; Rainbow^fl/+^*, we tracked clones over a diverse set of time windows, spanning birth to P60 (**Figure 5C**). We reasoned that the accrual of cells in clones between time of induction and time of harvest could reveal clonal expansion characteristics within discrete time windows. Tamoxifen dose was optimized to induce sparse labeling of cells, which then divide to form monochromatic clones (mCerulean, mCherry, or mOrange), enabling robust visualization and clone extraction (**Figure S5B**). A custom computational pipeline—combining machine learning and analytical approaches—was developed to reliably extract clones from the endothelium for a suite of downstream analysis, including spatial coordinates, arrangement, and size of individual clones (**Figure 5D, STAR Methods**). We focused our analysis on two colors, mCherry and mOrange, since mCerulean was disproportionately overrepresented, rendering clone calling difficult in the mCerulean channel (**Figure S5J**). The statistics of clone tagging were similar for mCherry and mOrange (**Figure S5C-F**), showing internal robustness of this approach.

We first measured the orientation of each clone relative to the direction of flow (**Figure 5E**) and found them to predominantly align with the direction of flow across all inducible time windows (**Figure 5F**). There were occasional clones with branching that contributed to circumferential growth (**Figure 5F**). This result demonstrates that mitotic orientation at individual cell-level, measured through spindle angles, is also captured at the scale of clones. We next asked whether larger clones, i.e., clones that had proliferated more, tended to be closer to other larger clones (and vice versa), or if they were stochastically interspersed with smaller or non-dividing clones. To assess the possibilities, we created three “simulated” endothelium from the collection of clone sizes, with (1) randomly distributed clones, (2) clones with biased positioning in the center of the tissue, and (3) clones with biased positioning on the sides of the tissue (**STAR Methods**). To all clones in the real and simulated data, a centroid position was assigned followed by nearest neighbor analysis (**Figure 5G, STAR Methods**). We found that the spatial coordinates of experimental clones were indistinguishable from those randomly distributed, irrespective of clone size (**Figure 5H)**. The results on clone size and distributions were consistent across different channels (**Figure S5M-O**).

To assess whether clonal characteristics varied across the two proliferative peaks, both average clone size as well as the largest clone size were measured and found to be significantly higher during the first wave than the second (average clone size for first wave is 2.69, average clone size for second wave is 2.31, p < 10^-8^) (**Figure 5I-J, S5K,L,P-R**). Consistently, the clone size from inducible windows spanning both peaks (P5-P60) had the highest average size (**Figure 5I-J**). The difference in clone size between the first and second wave was also validated using the 15-combination Mosaic system ^20^ (**Figure S5G-I**).

Finally, we computed the Gini coefficient, a measure of inequality within a population, for clone size distributions during each time window. The Gini coefficient is constrained between 0 and 1 and a higher value corresponds to a more unequal population, which in our context refers to higher variability in clones’ size within a window (**Figure 6A**). We found that the clones were “more equal” for the second wave compared to the first, as the average Gini coefficient of second wave clones was 0.19 compared to the average Gini coefficient of 0.27 in the first wave (**Figure 6B, S6A-E**). Consistently, the Gini coefficient for clone distributions from time windows spanning both waves was in between the two extremes (**Figure 6C**). These results collectively demonstrate that clonal characteristics, including the clone size and the population variation thereof, are distinct across the two proliferative waves.

**Figure 6.**
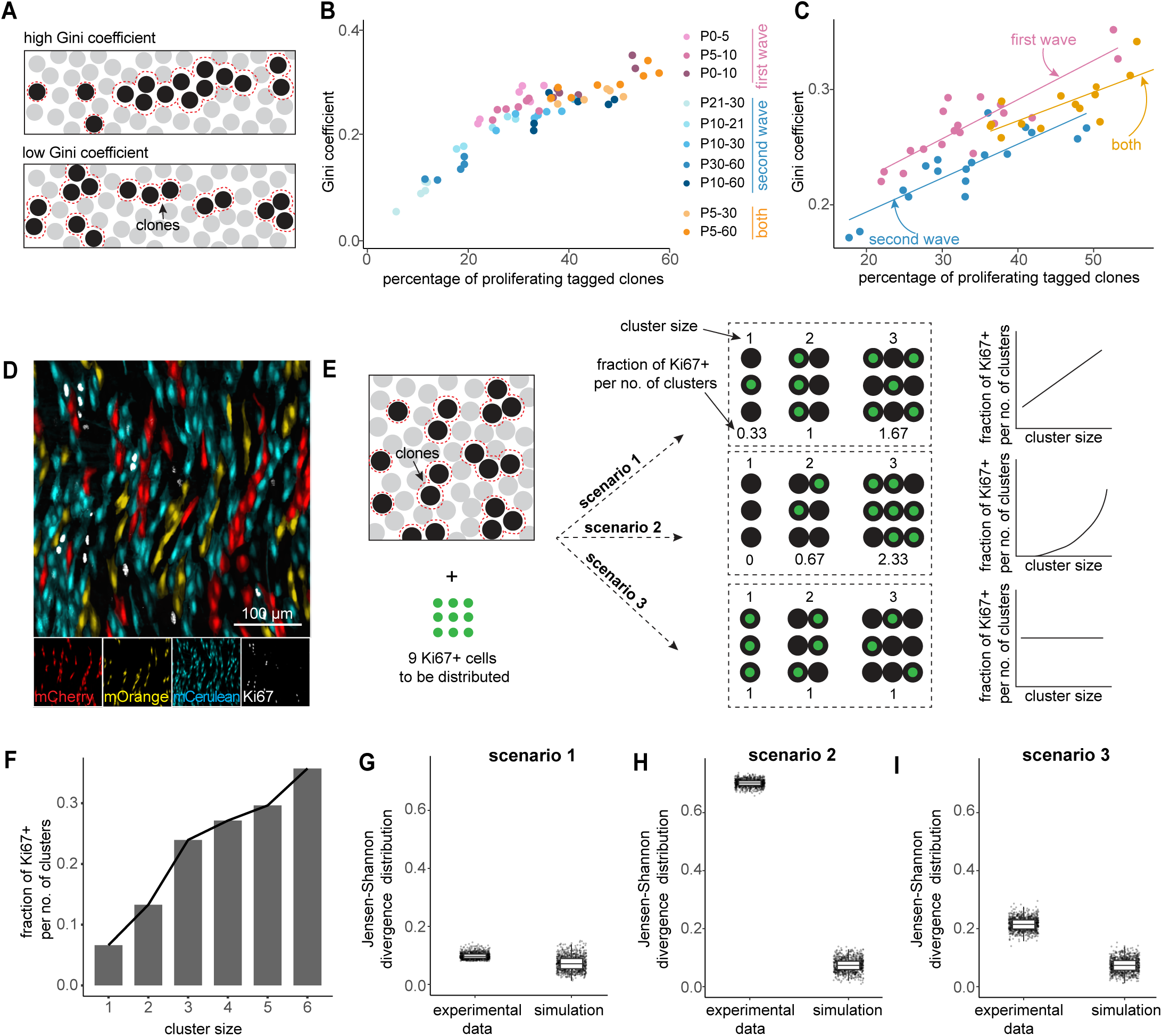
Clonal outgrowth occurs through cell-autonomous mechanisms. (A) Schematic of scenarios of clones that would either result in a high or low Gini coefficient. (B) Gini coefficient of each time window, color-coded by whether it is found within the first wave, second wave, or both. N = 5-8 animals per time point. (C) Gini coefficients of cluster sizes, calculated for different time windows, are categorized based on their association with the first wave, second wave, or both, and a regression line is fitted to each group. (D) Representative image of *en face* whole mount inducible rainbow aortas combined with Ki67 immuno-staining. Scale bar = 100 µm. (E) Schematic of the different scenarios of clonal behavior; scenario 1: clones do not have proliferative memory; scenario 2: clones have proliferative memory; scenario 3: proliferation is distributed equally amongst all clones. (F) The fraction of Ki67+ cells per number of clusters, against the cluster size, for experimental data within the first wave (P0-7). N = 3 animals. (G) Jensen Shannon divergence for experimental data and simulations for scenario 1 (clones do not have proliferative memory). (H) Jensen Shannon divergence for experimental data and simulations for scenario 2 (clones have proliferative memory). (I) Jensen Shannon divergence for experimental data and simulations for scenario 3 (proliferation is distributed equally amongst all clones).

### Clonal outgrowth occurs through cell-autonomous mechanisms

We sought to assess the mechanisms underpinning observed differences in clone sizes. One possibility is that the propensity to divide is memoryless or, mathematically speaking, Markovian at the cellular level (scenario 1). Another alternative is that clones exhibit proliferative memory, such that fast-dividing clones continue to do so and vice versa. This would imply that the proliferative potential is clone-intrinsic (scenario 2). Although unlikely, we cannot formally rule out yet another possibility that cells primed to undergo division must be distributed uniformly to the clones, regardless of the clone size, such as through a clone-size sensing negative feedback (scenario 3). To discriminate between these scenarios, we performed a new set of experiments simultaneously measuring clonal distributions using the Rainbow system coupled with Ki67 staining to identify actively dividing cells (**Figure 6D**). We reasoned that if the clones exhibited proliferative memory, the fraction of Ki67+ cells per number of cells in a clone would be higher for larger clones. Conversely, if no clonal memory were to be present, the number of Ki67+ cells per clone would scale perfectly with clone size (**Figure 6E**). We found the latter to be the case, where the fraction of Ki67+ cells scaled linearly with increasing clone size (**Figure 6F**). We quantitatively confirmed our observations by simulating the three scenarios raised above and found the Jensen-Shannon (**Figure 6G-I**), Kullback–Leibler divergence (**Figure S6F**), and Wasserstein distances (**Figure S6G**) between the simulated and experimental distributions to be most consistent with the memoryless scenario. We further probed whether the larger clones from the first wave continued to preferentially divide in the second wave. Here we induced clone tracing with tamoxifen in the first wave and measured Ki67+ cells during the peak of the second wave. We found no clear preference of Ki67+ cells for larger clones (**Figure S6H-J**). Taken together, the shape of observed clonal distributions is likely driven by individual cells’ propensity to divide, akin to a coin toss for each round of cell division.

### Endothelial cell cycle *in vivo* is remarkably short

Our observations on clonal sizes and distributions thereof led us to ask how the clonal phylogenies differed across the two waves and whether cell cycle durations could be extracted. We sought to experimentally answer these questions by performing *in vivo* cellular labeling with EdU during the first and second proliferative waves. At time of injection, EdU becomes incorporated into the newly synthesized DNA in S phase cells, and the label is passed on stoichiometrically to daughter cells upon cell division. As such, the amount of EdU in a daughter or granddaughter cell can be harnessed to infer how many generations of cell division had occurred since the time of injection. As EdU is non-discriminately incorporated into any cell in S phase, we inadvertently also labeled dividing smooth muscle cells. However, the developed computational pipeline, using a combination of endothelial nuclear signal (ERG) and nuclear orientation (smooth muscle cells orient perpendicularly to endothelial cells), was able to reliably segment and retain EdU signal from endothelial cells only (**Figure 7A**). By defining a clone as a group of cells that had touching cell borders (PECAM) (**STAR Methods**), we were able to reconstruct a lineage tree for each clone. With these trees established, we compared the two waves over the same fixed duration of 24 hours and focused on maximum clone size to understand the shortest possible cell cycle duration. Similar to the inducible rainbow clonal results—albeit over much shorter windows and thus with greater accuracy—the EdU clones were significantly larger in the first wave, with a maximum clone size of 12 cells (**Figure 7B**). In contrast, the largest clones observed in the second wave consisted of 4 cells (**Figure 7C**). We confirmed that circulating EdU has a short half-life and is no longer active after ∼60 min, ensuring cells are not progressively labeled and further consolidating the validity of our approach (**Figure S7A-B, STAR Methods**).

**Figure 7.**
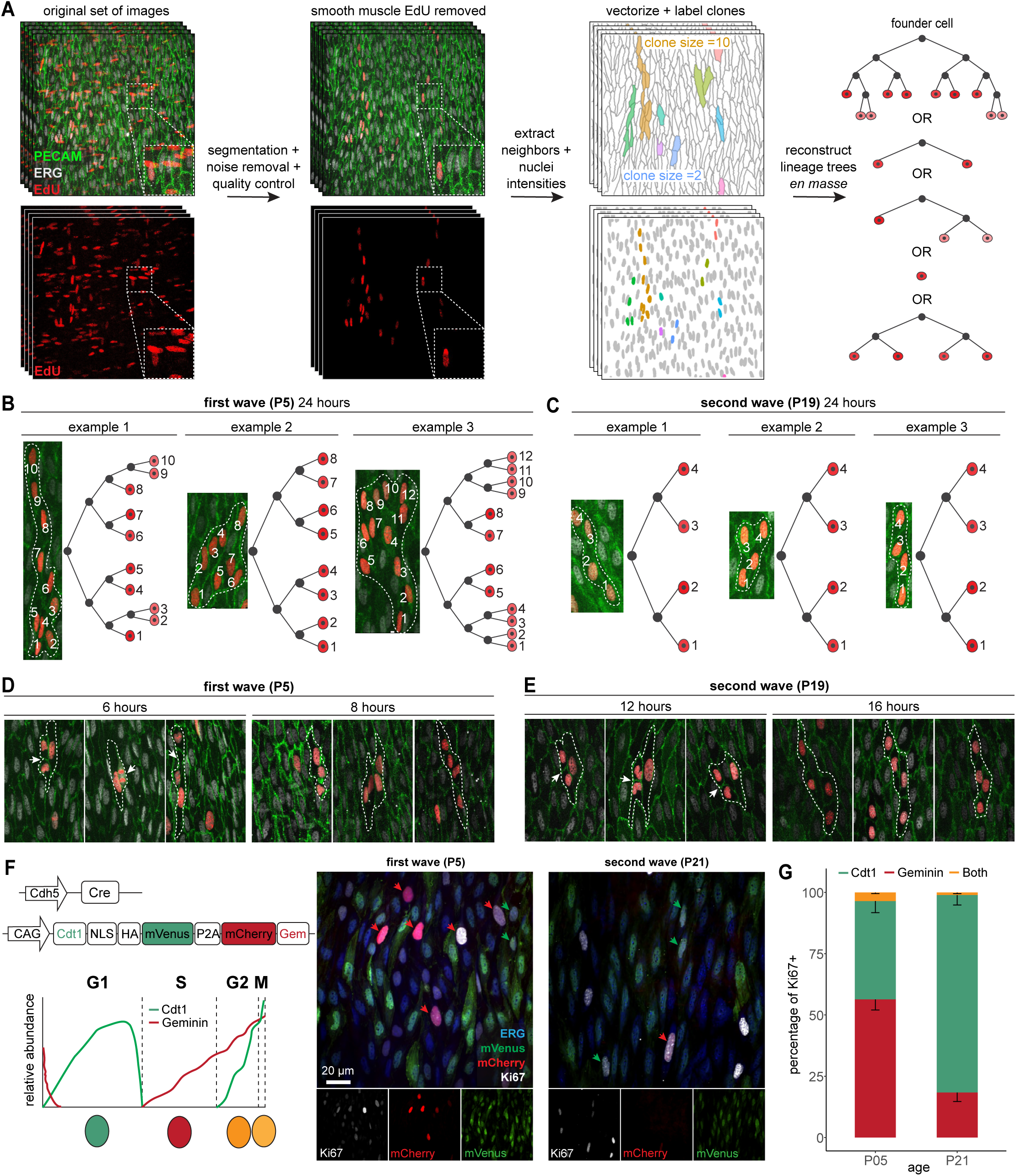
Endothelial cell cycle in vivo is remarkably short. (A) Work flow of *en face* whole mount image processing through machine learning segmentation to filter out smooth muscle cell signals and identify endothelial EdU clones. Signal intensity of each EdU+ cell is then used to reconstruct lineage trees. (B) Largest clones and their corresponding reconstructed lineage trees from 24 hours of EdU pulse-chase at P5 (first wave). Clones identified from N = 3 animals. (C) Largest clones and their corresponding reconstructed lineage trees from 24 hours of EdU pulse-chase at P19 (second wave). Clones identified from N = 3 animals. (D) Representative images showing recently divided EdU clones, 6 hours and 8 hours after EdU labeling at P5 (first wave). Dashed lines outline the identified clones and white arrows point to cells that have recently completed mitosis. (E) Representative images showing recently divided EdU clones, 12 hours and 16 hours after EdU labeling at P19 (second wave). Dashed lines outline the identified clones and white arrows point to cells that have recently completed mitosis. (F) Schematic of PIP-FUCCI construct, driven by tamoxifen-inducible Cdh5-Cre expression. Representative images of *en face* whole mount images from first wave (P5) and second wave (P21) *Cdh5-Cre; PIP-FUCCI^fl/fl^* animals. Red arrows point to cells that are Ki67+ and mCherry+, and green arrows point to cells that are Ki67+ and mVenus+. Scale bar = 20 µm. (G) Quantification of percentage of Cdt1 positive, Geminin positive, or double positive cells on the basis of Ki67+ cells, from first (P5) and second (P21) wave. Error bars represent standard deviation.

We next progressively reduced the induction-harvest time windows for each wave, ranging from 6 to 16 hours (**Figure 7D**). Remarkably, we observed clone sizes of three cells or higher even with short 6 and 8 hour time windows for the first wave (**Figure 7D**). To conservatively estimate the cell cycle duration, we focused on clone sizes of three instead of two since it is possible that the initial cell was only labeled with EdU at the end of its S phase. Remarkably, during the first wave, a 6 hour induction-harvest window showed several clusters of three cells, with two of the cells having recently completed cell division as indicated by post-mitotic nuclear morphology (**Figure 7D, white arrows**). Accordingly, by 8 hours of EdU labeling, we observed completed cell cycles, with observed clusters of three and four cells (**Figure 7D)**. Thus, we found that differentiated endothelial cells undergo cell division with a cell cycle duration as short as 6 hours in the first wave, significantly faster than previously estimated for endothelial cells in other vascular beds ^21,22^. This timescale resembles cell cycle durations associated with early embryonic stages ^23,24^. We next performed the same analysis for the second wave and observed the cell division duration to be approximately 12 hours, double that of the first wave (**Figure 7E**). Together, the quantitative approach applied revealed that endothelial cells can undergo remarkably rapid cell division, with each wave operating at different cell cycle duration timescales.

Given the stark difference in cell cycle durations between the first and second wave, as well as the changes in cell size, we asked whether a particular stage of the cell cycle was affected. Specifically, we wondered whether G1 was reduced in the first wave to shorten total cell cycle duration and produce smaller cells, as is known to occur in other systems ^25^. To test these possibilities, we used an *in vivo* model of PIP-FUCCI (*Cdh5 Cre; PIP-FU*CCI*^fl/fl^*) in combination with Ki67 immuno-staining. Ki67+ cells were scored as being in G1 (Cdt1+ and Geminin-), S Geminin+), and G2/M (Cdt1+ and Geminin+) (**Figure 7F**). We found that the ratio of cells in G1 was 2-fold higher for the second wave (P21) as compared to the first wave, consistent with a shortened G1 in the first wave (**Figure 7G**).

### Cellular extrusion is a critical regulator of tissue scaling during vascular growth

With the cellular parameters established (endothelial cell number, clonal dynamics, and cell cycle duration), we questioned how well would they adjust to the total tissue (aorta) scale. We assessed this by developing a stochastic mathematical model (**Figure 8A, STAR Methods**) consisting of two waves of cell division (*k*)—the first wave beginning at postnatal day 1 and the second wave starting at *T_0_* days after the beginning of the first wave. Each wave was characterized by *C_k_* rounds of division, and *f_k_* fraction of existing cells undergo division. The duration of cell cycle was sampled from an exponential distribution with a mean *T_k_*. To keep the model relatively general, we also included an effective rate of cell extrusion *P_k_*, which is observed in other monolayer systems such as the epithelia ^26^. The probability of a daughter cell to be lost after division is denoted by *P_k_*; therefore, on average, each cell division results in 2 - *P_k_* cells. We used the Markov Chain Monte Carlo (MCMC) algorithm ^27^ to identify the parameters that recapitulate the experimentally observed data (**Figure 8B, S8A-E, Table S3**). To test the effect of removing extrusion, we adopted two approaches: first, by keeping all other parameters constant and setting extrusion to 0 (*P_k_* = 0) (**Figure 8C**); second, by running a new set of MCMC and forcing extrusion to be 0 (**Figure 8E**). The former approach overestimated the total cells by an average of 1.75-fold (range of fold change: 1.38-2.33) (**Figure 8C,D**), while the latter resorted to 2-fold increase in the effective cell cycle durations than what was experimentally observed (**Figure 8E**) to compensate for the loss of cell extrusion.

**Figure 8.**
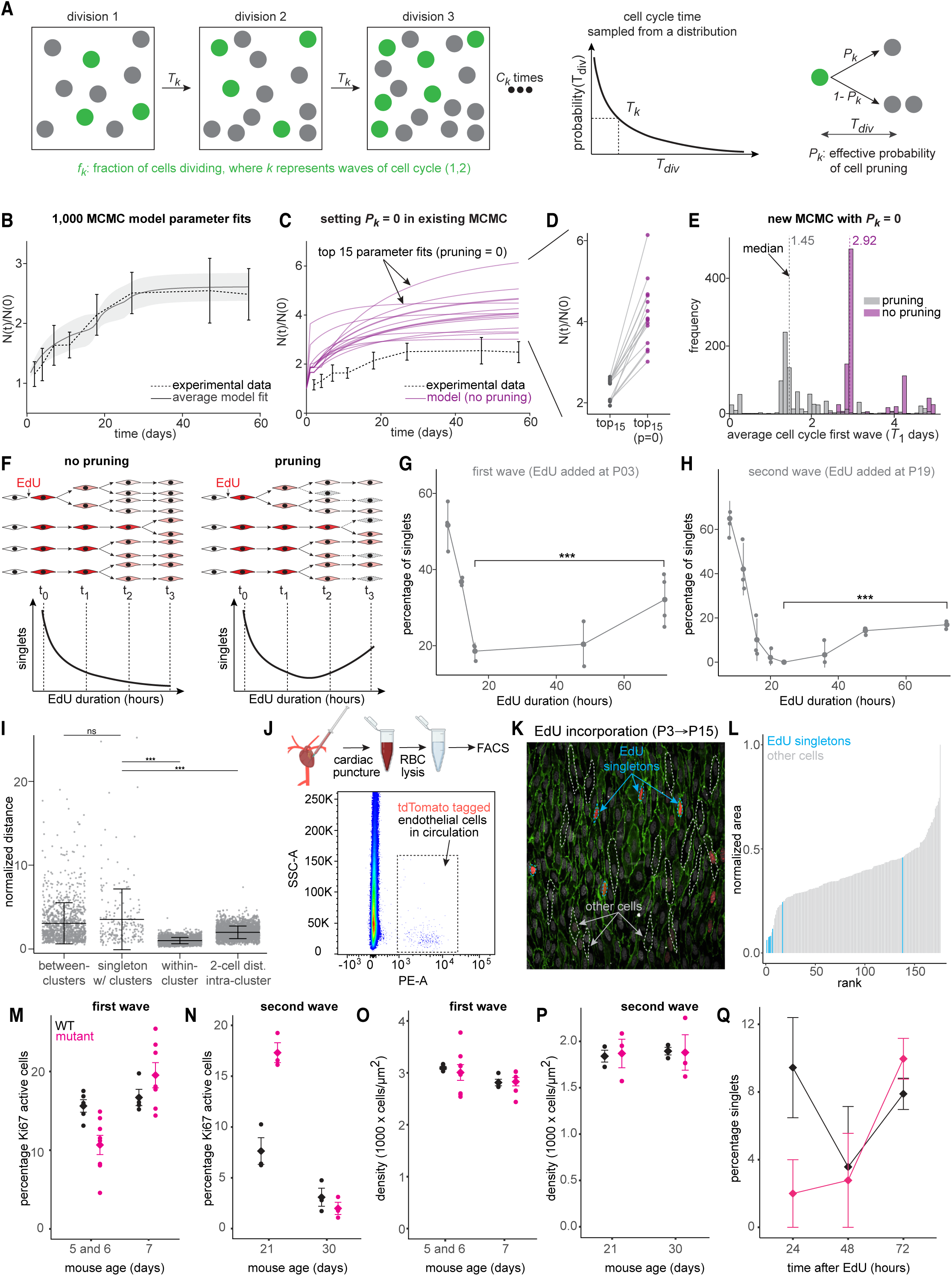
Cellular extrusion is a critical regulator of tissue scaling during endothelium expansion and growth. (A) Schematic showing key parameters from the mathematical model. There are two waves of cell division and each wave consists of *C_k_* rounds of division. In each round, *f_k_* fraction of cells go through division. The duration of cell cycle, *T_k_*, is sampled from an exponential distribution. The effective rate of extrusion (or pruning) is *P_k_* and each round of cell division results in 2-*P_k_* cells. (B) Top 1000 Markov Chain Monte Carlo (MCMC) model fits (solid line) compared to the real data for number of cells (dashed line). The error bar of real data is shaded and the error bar of the model is in lines; error bars represent standard deviation. (C) Top 15 Markov Chain Monte Carlo (MCMC) model fits with *P_k_*=0 compared to the real data, for number of cells. (D) Pairwise comparison between the top 15 Markov Chain Monte Carlo (MCMC) models with *P_k_* =/ 0 (gray) and pk = 0 (purple). (E) Distribution of average cell cycle of first wave in Markov Chain Monte Carlo (MCMC) model with pk =/ 0 (gray) and pk = 0 (purple). (F) Schematic of expected percentage of EdU singlets across time without or with cell loss. (G) Experimental data of percentage EdU clone singlets during first wave (EdU pulse at P3). Each dot represents a biological replicate and the error bar represents standard deviation. ***p<0.05 using independent t-test. N = 3-4 animals per group. (H) Experimental data of percentage EdU clone singlets during second wave (EdU pulse at P19). Each dot represents a biological replicate and the error bar represents standard deviation. ***p<0.05 using independent t-test. N = 2-3 animals per group (I) The normalized distance between all clusters, between singletons and clusters, between cells within a cluster, and between cells that are 2 cells apart within a cluster. The horizontal line indicates the mean and error bars represent standard deviation. p < 0.001 Welch’s t-test. (J) FACS analysis of whole blood to detect tdTomato-tagged endothelial cells in circulation, from P7 animals. N = 5 animals combined into one experiment. (K) Representative image of EdU singlets after multiple days of chase (image shown is EdU injected at P3 and harvested at P15). Dashed lines outline EdU singlets and other cells. (red: EdU; white: ERG; green: PECAM). (L) Distribution of normalized surface area of 176 cells from 1 aorta, aged P3-15. EdU singlets are marked in blue. (M) Percentage Ki67 active cells during the first wave from wildtype (black) and *Osx-Cre; Fgfr1^fl/fl^ Fgfr2^fl/fl^* (pink) animals. The diamond indicates the mean and error bars represent standard error of the mean. N = 4-8 animals per group. (N) Percentage Ki67 active cells during the second wave from wildtype (black) and *Osx-Cre; Fgfr1^fl/fl^ Fgfr2^fl/fl^* (pink) animals. The diamond indicates the mean and error bars represent standard error of the mean. N = 3 animals per group. (O) Endothelial density in the aorta during the first wave from wildtype (black) and *Osx-Cre; Fgfr1^fl/fl^ Fgfr2^fl/fl^* (pink) animals. The diamond indicates the mean and error bars represent standard error of the mean. N = 4-8 animals per group. (P) Endothelial density in the aorta during the second wave from wildtype (black) and *Osx-Cre; Fgfr1^fl/fl^ Fgfr2^fl/fl^* (pink) animals. The diamond indicates the mean and error bars represent standard error of the mean. N = 3 animals per group. (Q) Percentage of EdU singlets after 24, 48, and 72 hours of EdU labeling, from P19 wildtype (black) and Osx-Cre; Fgfr1^fl/fl^ Fgfr2^fl/fl^ (pink) animals. The diamond indicates the mean, connected by a trendline, and error bars represent standard error of the mean. All 24, 48 hour images were acquired using a 60X objective and all 72 hour images were acquired using a 20X objective. N = 3-5 animals per group.

We next sought to probe the presence of cell loss experimentally. We reasoned that in the absence of cell extrusion, the fraction of EdU+ singlets, i.e., cells which have not divided yet, should decrease monotonically over time as cells divide and become doublets and triplets (**Figure 8F, left**). Conversely, if extrusion were to be prevalent, we would expect the fraction of EdU+ singlets to first decrease and subsequently increase over time as EdU+ doublets and triplets lose cells to become singlets again (**Figure 8F, right**). Our experimental findings for both waves were consistent with the latter model, where the fraction of EdU+ singlets initially decreases before increasing again (**Figure 8G,H, S8F**). We ruled out the possibility that new singlets were being continuously labeled with EdU circulating in the blood through two arguments: one, EdU has a short distribution half-life of 1.4 minutes in mice ^28^, and two, our experiments revealed that there was no pharmacologically active circulating EdU beyond 1 hour of the initial injection (**Figure S7A-B**). To further ensure that we were not erroneously identifying daughter cells that had separated as singlets, we compared the distance between clusters (clone size >1) and clusters, singlets and clusters, and cells within the same cluster, and found that the pairwise distance between singlets and clusters was indistinguishable from that of clusters and clusters (**Figure 8I, S8G-H**). We further confirmed the presence of concurrent cell extrusion during periods of high proliferation by performing FACS on whole blood and identifying endothelial cells that had entered circulation (**Figure 8J**). For the same volume of whole blood collected, P7 animals had significantly higher instances of endothelial cells in circulation than adults (**Figure S8L-P**). Furthermore, through confocal microscopy, we observed instances of endothelial cells actively extruding above the z-plane of the monolayer (**Figure S8I-J**). Notably, the singlets monitored over a longer time period (P3→P15) tended to be smaller cells, perhaps suggesting that they were also in the process of being lost from the monolayer of endothelial cells (**Figure 8K-L, S8K**). Together, these results highlight the remarkable ability of endothelial cells to coordinate cell division, growth, and extrusion over time to accomplish global aorta growth characteristics.

With the myriad of quantitative information at hand regarding normal growth of an aorta, we next asked which parameters were affected in a disease model of achondroplasia, where the aorta is shorter. Remarkably, we found that while the two waves of proliferation were retained in the achondroplasia mice, the second wave’s proliferative peak was significantly higher in mutants than in wildtype, before reaching baseline at P30 (**Figure 8M-N**). This increase in proliferation despite the smaller aorta size led us to check for changes in cell density and cell loss, arguing that at least one of these two parameters had to be altered to accommodate for the increase in proliferation. We did not find any difference in cell density between wildtype and mutant conditions (**Figure 8O-P**). However, cell loss or extrusion appears to be increased in the mutants; while the percentage of EdU positive singlets did not change significantly between 24, 48, and 72 hours in the control, percentage EdU positive singlets increased significantly between 48 to 72 hours in the mutant after an initial decrease. (**Figure 8Q**). Therefore, the integration of cell proliferation, density, and extrusion collectively determine the final size of an endothelium.

## DISCUSSION

In most organs, postnatal growth is accomplished by the proliferation of tissue-resident stem cells ^2^. Frequently, these cells are confined to specialized niches and are also responsible for the homeostatic renewal of adult tissues ^2,3^. A clear exception is the liver, where postnatal growth and homeostatic renewal are both accomplished by differentiated hepatocytes ^2^. The endothelium of the aorta also follows this pattern. In fact, our findings from clone tracing and transcriptomic analyses indicated that the growth of the aortic endothelium is driven by the local proliferation of differentiated cells fully committed to the arterial fate. This contrasts the current view that arterial fate is incompatible with proliferation ^10^. Importantly, aortic endothelial cells retain their contacts, as indicated by clonal lineage tracing, and if they do migrate, they do so collectively.

We found that in the aorta, mitotic proliferation of endothelial cells is stochastic and monotonic with equivalent opportunity for cells across the entire monolayer to undergo cell division. Consistently, proliferation at the clonal level was also Markovian or memoryless, as opposed to other clonal modes of growth reported in the literature, including pre-existing propensities for dominance and spontaneous dominance (akin to forest fires) ^29–32^. Interestingly, cell division follows the direction of blood flow, contributing to a 2.5-fold tissue expansion in overall length. Growth in diameter is fulfilled by a slight deviation of cell division angle (+/- 30°) from laminar flow in addition to increases in cell size. Mitotic orientation yields long clones, which are also aligned in parallel to the flow, and it is present at all time points examined until P30. The findings are also consistent with the cessation of diameter expansion after this time. We also observed a significant increase in cell size after P10, which aligns well with the change in the metabolic expression profile of endothelial cells.

Interestingly, our data and mathematical model revealed that the proliferative kinetics supersede the required number of cells by 1.75-fold. This paradigm has been documented in the nervous tissue, where the generation of neurons also exceeds the number of needed cells ^33^. In nervous tissue, cell number adjustment relies on downstream synapses with target cells; unsuccessful connections or lack of stimulation lead to the elimination of the neuron by apoptosis ^33,34^. While we do not fully understand the mechanisms associated with endothelial cell extrusion, we could detect circulating endothelial cells at times when proliferation kinetics are higher. The implication is that a large number of endothelial cells must be lost at times when large vessels are growing. In fact, this paradigm aligns well with the presence of circulating endothelial cells in cord blood ^35^ and in other scenarios associated with high endothelial cell growth. This notion implies that circulating endothelial cells are likely exiting vessels as extruded and viable cells. Recent elegant studies on skin capillary networks found active post-natal remodeling accomplished by loss of cells with regression of the plexus and rearrangement of endothelial cells ^36^.

During the course of our analyses, we also found that the kinetics of proliferation was biphasic, with two waves interrupted by a relatively quiescent period. The first wave showed an asynchronous pattern and was extremely fast (6h or less), as shown by EdU kinetics and PIP-FUCCI evaluations. In contrast, the second wave was synchronous with slower cell cycle kinetics (12h). Short cell cycle times have been reported during gastrulation and in some regions of embryonic ectoderm (2.2-3.5h) ^37–39^. However, comparably short cell cycle durations have not been found postnatally nor in adult tissue-stem cells ^23,40^. Interestingly, in the case of hepatocytes, rapid regeneration has been attributed to their status of ploidy ^41^.

While the underlying mechanisms that regulate each of the mitotic waves are unclear, the consistency and reproducibility of these proliferation patterns suggest that they may be hard-wired and controlled by cell-intrinsic mechanisms. In fact, by including genetic perturbations that altered the vertebral column length, we found that the temporal occurrence of two waves and the quiescent period were unaltered. Our observations imply that the final size adjustment of the dorsal aorta and scaling to the vertebral column is mediated by endothelial cell loss into the circulation, as implied by the evaluation of EdU+ singlets in the achondroplasia model. In sum, our study underscores the power of quantitative approaches in revealing several unexpected and hitherto unappreciated features regulating aorta growth and scaling. This includes identification of discontiguous proliferative waves, fast cell cycle kinetics of resident differentiated endothelial cells, and extrusion.

A key finding from our studies was the drastic suppression of cell proliferation by P10 (quiescent period), which marks the initiation of dramatic elongation of the aorta and the vertebral column. At this time, endothelial cells were extremely packed and the tissue showed the highest density than any of the other time points, creating a mechanical stress anisotropy ^42,43^. This control strategy of rapid cell divisions without an increase in cell size early on appears to serve as a “reservoir” of cells (akin to early embryogenesis ^44–48^) that can enlarge and specialize over time to meet the demands of rapidly growing aorta after P10. Curiously, the dip in endothelial cell proliferation is also concurrent with a spike in mean wall shear stress and changes in the tunica media ^49^. Whether these changes reflect any direct relationship is, at this point, unclear.

The relationship between cell size reduction and suppression of proliferation is also another interesting causational posibility. Experiments by Watt ^50^ using adhesive micropatterns to control cell geometry showed that placing epidermal stem cells on small micropatterns decreased their DNA synthesis and induced differentiation. In contrast, plating these cells in larger micropatterns reduced differentiation. The implication was that density provided the efficient tissue-autonomous regulation of self-renewal and differentiation. In more recent work also in epithelium, spatiotemporal control of cell proliferation was found to provide feedback information imparted by the degree of tissue crowding and local changes in force distribution ^51^. Importantly, the effect of mechanical confinement on cell function is consistent with previous work linking tension to changes in chromatin organization ^52,53^. Interestingly, we found that repair of an aortic wound following mechanical injury resulted in robust and local endothelial proliferation, which did not stop after wound closure, yielding a significant hyper-density state. This aberrant cell packing is resolved by 60 days when the tissue gradually reverts to its previous cell density ^17^. At that time, the mechanism was unclear; now, given the present results, we might consider this return to normal density to be mediated by endothelial cell extrusion. Naturally, additional experiments are required to fully investigate processes and mechanisms.

## LIMITATIONS OF THE STUDY

Our findings indicate that the postnatal expansion of the aorta is fully reliant on the proliferation of differentiated resident arterial cells, and the aorta scales to fit deviations in both length and disease states. Admittedly, our studies were confined to endothelial cells (tunica intima) and excluded two adjacent important tissues: vascular smooth muscle cells (tunica media) and an external layer of connective tissue (tunica adventitia). While we noted the robust proliferation of smooth muscle cells (EdU uptake and Ki67 staining), we have refrained from evaluating cells other than the endothelium to focus the study and gain depth first in the endothelium. In addition, and as previously mentioned, the mechanisms that trigger the two waves of proliferative activity and the hiatus of proliferation at P10 are extremely intriguing and require additional attention. Finally, the mechanisms associated with endothelial cell extrusion should be addressed in future studies at a greater depth.

## RESOURCE AVAILABILITY

### Lead contact

Further information and requests for resources and reagents should be directed to and will be fulfilled by the lead contact, M. Luisa Iruela-Arispe.

### Materials availability

No new reagents or material were generated in this study.

### Data and code availability

All data generated in this manuscript has been deposited at: https://drive.google.com/drive/folders/1LVv6ynssGyE1rvt0NK81DhfL-Wl5Nna8?usp=sharing

All code in this manuscript has been deposited at: https://github.com/GoyalLab/aorta_growth_PiBraun_etal/

## ACKNOWLEDGEMENTS

We thank members of the Arispe and Goyal labs for helpful discussions and comments on the manuscript. We especially thank Dr. A McDonald, who initiated this project while a graduate student at UCLA in the Arispe lab. We thank Sam Buchanan and Snezana Mirkov from the Arispe lab for their assistance with animal husbandry and Emanuelle Grody from the Goyal lab for her assistance with sequencing single-cell RNA sequencing libraries. We thank Madeline Melzer, Ian Mellis, and Nitu Kumari from the Goyal lab for discussions on the experimental design and computational analyses. We thank the NUSeq Core, Flow Cytometry Core, Feinberg Information Technology, Northwestern University Information Technology, Quest High Performance Computing Cluster, and Center for Advanced Microscopy and Nikon Imaging Center at Northwestern University Feinberg School of Medicine for their services. AM acknowledges funding from American Heart Association (AHA) 23POST1022462. VLB acknowledges funding from National Institutes of Health (NIH) R35 HL139950. YG acknowledges support from Northwestern University’s startup, AHA Collaborative Science Award (24CSA1256987 to MLIA and YG) and Burroughs Wellcome Fund Career Awards at the Scientific Interface. JB, SD, and SC were supported by the Burroughs Wellcome Fund Career Awards at the Scientific Interface award and to YG. JB and YG also acknowledge DAAD IFI Fellowship for supporting this work. SD and YG also acknowledge Seed funding for Collaboration and Partnership Projects (SCPP) from IIT Bombay IRCC vide project no RD/0523-IOE00I0-213 for facilitating part of their collaborative research. YG is a CZ Biohub Investigator. MLIA acknowledges funding for this project from NIH R35HL140014 and R01HL178787.

## AUTHOR CONTRIBUTIONS

MLIA and YG conceived and designed the study. DP and JB designed and performed a majority of the experiments and analyses, respectively. SD formulated the mathematical model with input from YG. DP performed experiments related to *Osx-Cre, Fgfr1^fl/fl^ Fgfr2^fl/fl^* mice with input from DO. DP and PB performed experiments related to *Cdh5-Cre, PIP-FUCCI^fl/fl^* mice with input from VLB. AM prepared single cell sequencing libraries. FM, JB, and YG performed single-cell RNA sequencing analysis with input from MLIA and DP. SRS performed experiments related to microcomputed tomography. SC assisted in data annotation for machine learning image segmentation. LGO performed experiments related to *Cdh5-Cre, iMb2-Mosaic, iChr2-Mosaic* mice with input from RB. YG prepared a majority of the figures with some contributions from DP, and input from JB and MLIA. MLIA and YG wrote the manuscript with input from all authors.

## DECLARATION OF GENERATIVE AI IN THE WRITING PROCESS

The entire first draft was written without the use of any generative AI. To improve the wording of a sentence, the authors used Claude.ai, but such instances were rare. After using Claude.ai, the authors reviewed and edited the content as needed and take full responsibility for the content of the publication.

## SUPPLEMENTAL INFORMATION

**Table S1:** Vertebral level of each aorta landmark.

| Age_ID | Age | Sex | celiac trunk | intercostal 10 | intercostal 9 | intercostal 8 | intercostal 7 | intercostal 6 | intercostal 5 | intercostal 4 | intercostal 3 | intercostal 2 | intercostal 1 | L subclavian |
| --- | --- | --- | --- | --- | --- | --- | --- | --- | --- | --- | --- | --- | --- | --- |
| P3_11 | 3 | F | L1 | L1 | T13 | T12 | T12 | T11 | T9 | T9 | T7 | T6 | T5 | T2 |
| P3_1 | 3 | M | L1 | L1 | T13 | T12 | T12 | T11 | T10 | T8 | T7 | T6 | T5 | T2 |
| P3_2 | 3 | M | L2 | L1 | T13 | T13 | T12 | T11 | T10 | T9 | T8 | T6 | T5 | T1 |
| P5_1 | 5 | M | L1 | L1 | T13 | T12 | T11 | T10 | T9 | T8 | T7 | T6 | T5 | T3 |
| P5_2 | 5 | M | L2 | L1 | T13 | T12 | T11 | T11 | T10 | T9 | T7 | T6 | T5 | T2 |
| P5_14 | 5 | F | L1 | T13 | T12 | T11 | T10 | T10 | T8 | T7 | T6 | T5 | T4 | T1 |
| P5_13 | 5 | F | L1 | T13 | T13 | T12 | T11 | T10 | T9 | T8 | T7 | T6 | T4 | T2 |
| P7_11 | 7 | F | L1 | T13 | T12 | T11 | T10 | T9 | T9 | T8 | T6 | T6 | T5 | T2 |
| P7_12 | 7 | F | L1 | T13 | T12 | T11 | T11 | T9 | T8 | T7 | T6 | T6 | T5 | T2 |
| P7_1 | 7 | M | L1 | T13 | T12 | T11 | T10 | T9 | T8 | T7 | T6 | T5 | T5 | T2 |
| P7_2 | 7 | M | L1 | T13 | T12 | T11 | T11 | T9 | T8 | T7 | T6 | T6 | T5 | T2 |
| P10_2 | 10 | M | L2 | L1 | T13 | T12 | T11 | T10 | T9 | T8 | T7 | T6 | T5 | T2 |
| P10_3 | 10 | M | L1 | T13 | T12 | T11 | T10 | T9 | T8 | T7 | T6 | T5 | T5 | T2 |
| P10_11 | 10 | F | L2 | T13 | T12 | T12 | T10 | T9 | T8 | T7 | T6 | T5 | T5 | T1 |
| P10_12 | 10 | F | L1 | T13 | T12 | T12 | T10 | T10 | T9 | T7 | T6 | T5 | T5 | T2 |
| P14_11 | 14 | F | L2 | L1 | T13 | T12 | T11 | T10 | T10 | T8 | T8 | T7 | T6 | T2 |
| P14_12 | 14 | F | L1 | T13 | T12 | T11 | T11 | T9 | T8 | T7 | T6 | T5 | T5 | T2 |
| P21_13 | 21 | F | L2 | L1 | T13 | T12 | T12 | T11 | T10 | T9 | T8 | T7 | T6 | T3 |
| P21_14 | 21 | F | L1 | L1 | T12 | T12 | T11 | T10 | T9 | T8 | T7 | T6 | T5 | T2 |
| P21_3 | 21 | M | L1 | T13 | T12 | T11 | T10 | T9 | T9 | T7 | T7 | T6 | T5 | T2 |
| P21_4 | 21 | M | L2 | L1 | T13 | T13 | T12 | T11 | T10 | T9 | T8 | T7 | T6 | T3 |
| P30_11 | 30 | F | L2 | L1 | T13 | T13 | T12 | T11 | T10 | T9 | T8 | T7 | T6 | T2 |
| P30_12 | 30 | F | L2 | L1 | T13 | T12 | T11 | T11 | T9 | T9 | T7 | T7 | T6 | T2 |
| P30_4 | 30 | M | L1 | T13 | T12 | T12 | T11 | T10 | T9 | T8 | T7 | T6 | T5 | T2 |
| P30_3 | 30 | M | L2 | T13 | T12 | T11 | T11 | T10 | T9 | T8 | T7 | T6 | T5 | T2 |
| P60_11 | 60 | F | L3 | L1 | T13 | T12 | T12 | T11 | T10 | T9 | T8 | T7 | T6 | T3 |
| P60_12 | 60 | F | L2 | L1 | T13 | T13 | T12 | T11 | T10 | T9 | T8 | T7 | T6 | T3 |
| P60_2 | 60 | M | L1 | T13 | T12 | T11 | T11 | T10 | T9 | T8 | T7 | T6 | T5 | T2 |
| P60_4 | 60 | M | L2 | L1 | T13 | T12 | T11 | T10 | T9 | T8 | T7 | T6 | T5 | T2 |

**Table S2:** Different thresholds for Ki67 segmentation and resulting percentage Ki67+.

| Age | Aorta | Main threshold | 15% lower threshold | 15% higher threshold |
| --- | --- | --- | --- | --- |
| 3 | aorta1 | 0.201 | 0.228 | 0.179 |
| 3 | aorta2 | 0.162 | 0.183 | 0.144 |
| 5 | aorta1 | 0.123 | 0.141 | 0.110 |
| 5 | aorta2 | 0.141 | 0.162 | 0.120 |
| 5 | aorta3 | 0.122 | 0.146 | 0.101 |
| 7 | aorta1 | 0.128 | 0.146 | 0.112 |
| 7 | aorta2 | 0.157 | 0.179 | 0.134 |
| 7 | aorta3 | 0.152 | 0.172 | 0.135 |
| 10 | aorta1 | 0.021 | 0.027 | 0.017 |
| 10 | aorta2 | 0.026 | 0.031 | 0.022 |
| 10 | aorta3 | 0.050 | 0.059 | 0.044 |
| 14 | aorta1 | 0.053 | 0.060 | 0.048 |
| 14 | aorta2 | 0.053 | 0.059 | 0.047 |
| 21 | aorta1 | 0.103 | 0.114 | 0.093 |
| 21 | aorta2 | 0.135 | 0.149 | 0.123 |
| 21 | aorta3 | 0.159 | 0.177 | 0.142 |
| 30 | aorta1 | 0.037 | 0.044 | 0.033 |
| 30 | aorta2 | 0.070 | 0.080 | 0.062 |
| 30 | aorta3 | 0.026 | 0.031 | 0.022 |
| 50 | aorta1 | 0.049 | 0.055 | 0.044 |
| 50 | aorta2 | 0.024 | 0.030 | 0.020 |
| 50 | aorta3 | 0.019 | 0.025 | 0.015 |
| 60 | aorta1 | 0.015 | 0.017 | 0.013 |
| 60 | aorta2 | 0.010 | 0.012 | 0.008 |
| 60 | aorta3 | 0.013 | 0.015 | 0.011 |

**Table S3:** Symbols and parameters for mathematical model.

| symbol | parameter | Range of initial guesses | symbol | parameter | Range of initial guesses |
| --- | --- | --- | --- | --- | --- |
| $f_1$ | Fraction of existing cells dividing in wave 1 | (0, 0.5] | $T_1$ | Duration of cell cycle in wave 1 | [0.5, 2] days |
| $C_1$ | Rounds of cell division in wave 1 | [1, 10] (integer) | $P_1$ | Pruning rate in wave 1 | [0, 1] |
| $f_2$ | Fraction of existing cells dividing in wave 2 | (0, 0.5] | $T_2$ | Duration of cell cycle in wave 2 | [0.5, 10] days |
| $C_2$ | Rounds of cell division in wave 2 | [1, 10] (integer) | $P_2$ | Pruning rate in wave 2 | [0, 1] |
| $T_0$ | Time after the beginning of the first wave | [10, 25] days | | | |

**Video S1. Microcomputed tomography reconstruction of backbone and aorta**

## METHODS

### EXPERIMENTAL MODEL AND SUBJECT DETAILS

#### Animals

The following mouse strains were housed at Northwestern University: C57BL/6J, *Cdh5 Cre-ERT2; Rainbow^fl/+^*, and *Cdh5 Cre-ERT2; tdTomato^fl/fl^*. All animal procedures were performed in accordance with Northwestern University’s Institutional Animal Care & Use Committee. Mice were maintained in accordance with policies set forth by the American Association for Accreditation of Animal Care. C57BL/6J mice were purchased from Jackson Laboratories (JAX stock #000664). *Cdh5 Cre-ERT2; Rainbow^fl/+^* mice were generated by crossing *Cdh5 Cre-ERT2* mice (from Dr. Ralf Adams, available from Taconic Biosciences) with *Rainbow^fl/fl^* mice (from Dr. Irving Weissman ^54^). *Cdh5 Cre-ERT2; tdTomato^fl/fl^* were generated by crossing *Cdh5 Cre-ERT2* with *tdTomato^fl/fl^* mice (JAX stock #007914). These mice were maintained in Northwestern’s vivarium, in group housed cages and with food and water *ad libitum*. Genotyping was performed through Transnetyx. Where possible, an equal number of males and females have been used in experiments. The age at which the animals were used are indicated in the data. To induce recombination in *Cdh5 Cre-ERT2* mice at the appropriate ages, adults were intraperitoneally injected with a single 100 uL dose of 10 mg/mL tamoxifen dissolved in ethanol and sunflower seed oil. Pups younger than 21 days were orally gavaged with a single 5 uL dose of 20 mg/mL tamoxifen dissolved in ethanol and sunflower seed oil.

Double conditional *Fgfr1* and *Fgfr2* knockout mice (DCKO) (*OsxCre; Fgfr1^f/f^; Fgfr2^f/f^*) were generated by crossing *Osx-Cre* mice with double homozygous *Fgfr1* and *Fgfr2* floxed mice (DFF) as described previously ^15^. These mice were housed in a pathogen-free vivarium at Washington University in St. Louis where they were handled in accordance with standard use protocols, animal welfare regulations and the NIH Guide for the Care and Use of Laboratory Animals and approved by the Washington University Animal Studies Committee. Mice were group housed with food and water *ad libitum*. Where possible, an equal number of males and females have been used in experiments. The age at which mice were analyzed are indicated in the figure legends. Experimental details related to “Microcomputed tomography”, “Aorta dissection and en face immunofluorescence”, and “EdU in vivo labeling” that were performed on these mice are detailed elsewhere. Fixed backbones with aortas attached were shipped to Northwestern University in 50 mL falcon tubes in PBS and further processed.

*Cdh5 Cre-ERT2, iMb2-Mosaic; iChr2-Mosaic* mice were generated by crossing *Cdh5 Cre-ERT2* mice with *iMb2-Mosaic; iChr2-Mosaic* mice ^20^. These animals were housed at the Spanish National Cardiovascular Research Center vivarium, in group housed cages with food and water *ad libitum*. Genotyping methods for these mice have previously been described ^20^. Where possible, an equal number of males and females have been used in experiments. To induce recombination, animals were intragastrically injected with 60uL of 2 ug/uL of 4-hydroxytamoxifen (dissolved in ethanol and castor oil) at P5. The age at which animals were harvested is indicated in the data. After harvesting and fixing overnight in 2% PFA, backbones with aortas attached were shipped to Northwestern University in 50 mL falcon tubes in PBS and further processed.

*Cdh5 Cre-ERT2, PIP-FUCCI^fl/fl^* mice were generated by crossing *Cdh5 CreERT2* mice with PIP-FUCCI knock-in reporter mice ^21^. These animals were housed at the University of North Carolina, Chapel Hill vivarium, in group housed cages with food and water *ad libitum*, and animal experiments were approved by the University of North Carolina at Chapel Hill Institutional Care and Use Committee. Genotyping methods for these mice have previously been described ^21^. Where possible, an equal number of males and females were used in experiments. To induce recombination, 50 uL of 1 mg/mL tamoxifen dissolved in sunflower seed oil was injected intraperitoneally into animals on days P1-P3. After harvesting and fixing overnight in 2% PFA, backbones with aortas attached were shipped to Northwestern University in 50 mL falcon tubes in PBS and further processed.

## METHOD DETAILS

### Microcomputed tomography

Upon sacrifice, mice were perfused with 1X PBS through the left ventricle to remove blood in large vessels. To ensure adequate fill, aortas were ligated with 5-0 suture at the neck vessels and at the abdominal aorta distal to the renal arteries. Mice were then secondarily perfused with 1-2 mL of contrast agent (Vascupaint Ex Vivo Contrast Agent, MediLumine) to fill the aorta and branching vessels. Perfused samples were then allowed to cure at room temperature for up to 30 minutes before removal of all organs, leaving behind the backbone with aorta attached. These perfused samples were then stored in 2% PFA at 4°C until ready for scanning. Microcomputed tomography (microCT) imaging was performed using a Bruker Skyscan 1272 with 60 kVp tube voltage, 0.25 mm AI filter, and angular sampling with 0.4° rotation increments collected over 180° rotation. Three frames were collected for each projection direction, and a 20 pixel random detector motion was used to suppress ring artifacts.

### Measurement of vertebrae and aorta

Analysis of microCT data was performed in FIJI. To account for natural curvature of the spine and aorta, multiple points spaced out across the spine and aorta length were measured. To measure the length of the aorta, we noted the x, y, z positions of the left subclavian artery branch point, celiac trunk branch point, and thirteen points in between (ten intercostal arteries and three equally spaced positions between the left subclavian artery and first intercostal arteries). The distance D_1-2_ between adjacent measurement points x_1_,y_1_,z_1_ and x_2_,y_2_,z_2_ was determined using the quadratic formula D_1-2_ = { (x_1_ - x_2_)^2^ + (y_1_ - y_2_)^2^ + (z_1_ - z_2_)^2^ }^0.5^, then all distances were summed up to calculate the total length of the thoracic aorta. To measure the length of the spine, S1 was first identified based on its distinct morphology (**Figure S9**). Each vertebral centrum’s cranial x, y, z position was noted between L4 and T1; total length of the spine was then calculated in a similar manner as the aorta. Vessel circumference was measured by averaging the cross-section circumference across ten equally spaced points in the thoracic aorta.

### Identification of corresponding aorta landmarks and vertebral levels

The z position of each aorta landmark - including the left subclavian artery branch point, celiac trunk branch point, and right branching vessel of 10 pairs of intercostal arteries, were noted in FIJI. The z position of the cranial plane of each vertebra from C7 to S1 were also noted in FIJI (**Figure S9**). The vertebral level was assigned to each aorta landmark by determining if the z position of the vessel branch point fell between a certain vertebra’s cranial z position and the next lower vertebra’s cranial z position.

### Aorta dissection and en face immunofluorescence

Upon sacrifice by CO_2_ inhalation, mice were perfused with 2% PFA through the left ventricle. The length of the aorta, from left subclavian to celiac trunk, was measured before removal from the backbone. Under a dissecting microscope, adventitia and branching vessels were removed before the aorta was opened up longitudinally to expose the intima. Fileted aortas were pinned with the intima facing up in a 35 mm silicon-coated dish before additional fixation with 2% PFA overnight at 4C. After fixation, tissues were washed and incubated in Blocking/Permeabilization Buffer (0.3% TritonX-100, 0.05% Tween-20, 3% Normal Donkey Serum in 1X HBSS) for 1 hour at room temperature, then incubated with primary antibody solution (made up in Blocking/Permeabilization Buffer) overnight at 4 degrees on an orbital shaker. After washing, secondary antibody solution (made up in Blocking/Permeabilization Buffer) was applied for 1 hour at room temperature. To mount, aortas were stretched and pinned to its measured length on a new silicon-coated dish. A small glass coverslip was placed under the pinned aorta, and Prolong Gold Antifade Mountant (Invitrogen) was pipetted onto the tissue before another small glass coverslip was placed over the tissue to seal. After curing overnight at room temperature, the glass-tissue-glass sandwich was transferred onto a microscope slide for confocal microscopy.

### Confocal imaging

All confocal images were acquired using a Nikon A1R HD25 microscope. Multi-tile and Z-stack images were stitched and maximum intensity processed using NIS Elements software. Figure images were denoised using NIS Elements software.

### EdU *in vivo* labeling

Mice were intraperitoneally injected with EdU (dissolved in PBS, 100uL of 10mg/mL for mice older than P21, 50uL of 10mg/mL for pups). At time of harvest, animals were sacrificed and aortas were harvested and fixed. Click-iT^TM^ EdU Alexa Fluor 594 reaction was performed on aorta tissue according to manufacturer’s instructions (Invitrogen), before immuno-staining with primary and secondary antibodies as described previously. Mounted tissues were then imaged through confocal microscopy.

### Single cell RNA sequencing

Animals were sacrificed through CO_2_ inhalation before perfusion through the left ventricle with warm Versene solution (0.2g EDTA, 8g sodium chloride, 0.2g potassium chloride, 1.15g anhydrous sodium phosphate, 0.2g anhydrous potassium phosphate, dissolved in distilled water, pH 7.5) The aorta was dissected from the body and adventitia removed before being cut open to expose the endothelium. For the P5 and P10 timepoints, whole aortas were pooled into digestion solution (Liberase TH 70ug/mL, Sigma-Aldrich; DNase I, Qiagen; 10mM HEPES, Corning) and incubated at 37°C for 20 minutes while pipetting up and down every 2 minutes to generate a single-cell suspension. For P20 and P30 timepoints, aortas were cut open and pinned down with the endothelium facing up. After adding warm 1X trypsin (Gibco) for 5 minutes, a microfeather blade was used to gently scrape the endothelium of the aorta, as in previously established methods ^17^. The samples were incubated in 1X trypsin for an additional 5 minutes while pipetting up and down to generate a single-cell suspension. After neutralizing the digestion solution with DMEM containing 10% FBS, the cell suspension was filtered through a 20 uM cell strainer. Post-digestion cleanup was done prior to cell counting and viability assessment. Cell suspensions were loaded onto a Chromium Next GEM Chip G and run using the Chromium Controller iX (10X Genomics) to generate gel beads in emulsion (GEMs). Single-cell RNA sequencing libraries were constructed using the Chromium Next GEM Single Cell 3’ v3.1 (Dual Index). Sequencing was performed on a NextSeq2000 Sequencing System (Illumina). Gene expression matrices were generated by de-multiplexing, aligning, barcode processing, and UMI counting using CellRanger pipeline.

### Single cell RNA sequencing data analysis

The approach to perform single cell analysis, including quality control steps and doublet removal was inspired from our previous studies ^17,55–57^. The Cellranger output expression matrices were merged for each sample. The R package Seurat (v5.1.0) was used to cluster the cells in the merged matrix. Cells with less than 100 genes or more than 2e4 transcripts or 7.5% of mitochondrial expression were first filtered out as low-quality cells. The NormalizeData function was used to normalize the expression level for each cell with default parameters. The FindVariableFeatures function was used to select variable genes with default parameters. The ScaleData function was used to scale and center the counts in the dataset. Principal component analysis (PCA) was performed on the variable genes. The RunHarmony function from the Harmony package was applied to remove potential batch effect among individual samples.

Uniform Manifold Approximation and Projection (UMAP) dimensional reduction was performed using the RunUMAP function. The clusters were obtained using the FindNeighbors and FindClusters functions with the resolution set to 0.5. To ensure data quality, we first filtered out low-quality cell clusters by removing clusters with an average of fewer than 1,000 genes detected per cell (nFeature_RNA) and fewer than 2,000 total molecules detected per cell (nCount_RNA). Following this filtration, doublets were identified and removed. Doublets were identified by evaluating the expression of specific markers - Pecam1, Dcn, Prox1, Tagln, and Cd3e - and determining whether each marker exceeded a defined threshold. Threshold levels were set based on a histogram analysis of gene expression data. Cells expressing two or more markers above these thresholds were classified as doublets. Upon completing these quality control measures, we proceeded with data normalization and scaling, followed by the application of Harmony and UMAP, as described earlier. The cluster marker genes were found using the FindAllMarkers function. The cell types were annotated by overlapping the cluster markers with the published marker genes. Subsequently, we focused on isolating endothelial and proliferating cells by employing clusters that had been annotated using our previous mentioned methodology. For each subset of data, we repeated normalization, scaling, and the application of Harmony and UMAP to ensure consistency across analyses. Proliferating endothelial cells were identified by further clustering within the proliferating cell group and refining cell type annotations utilizing the aforementioned methodology. The dot plot was plotted using the DotPlot function. The violin plots were plotted using the VlnPlot function. The UMAP showing individual genes were plotted using the FeaturePlot function. Differential expression analysis between two group of cells was conducted using the FindMarkers function. Genes with adjusted p value smaller than 0.05 were considered significantly differentially expressed. The module scores were calculated based on genes in GO pathways downloaded from https://geneontology.org/.

### (Supplemental)#Measuring EdU bioavailability in mouse circulation

Adult mice were intraperitoneally injected with 100 uL of 10 mg/mL EdU dissolved in PBS, then sacrificed after 20 minutes, 1 hour, and 3 hours. Whole blood was obtained through cardiac puncture and centrifuged at 3000 rpm for 10 min for fractionation. Serum was collected and heat-inactivated at 56C for 30 min before storing at -20C. As a control, fresh EdU was mixed with fetal calf serum, then heat-inactivated and stored alongside collected mouse serum. Human dermal fibroblasts were serum starved overnight, then treated the next morning with media containing 10% of mouse serum or 10% EdU/fetal calf serum. After 3 hours, fibroblasts were fixed with 2% PFA before Click-iT^TM^ Alexa Fluor 594 reaction (Invitrogen). Fixed cells were then blocked, permeabilized, and immuno-stained with primary and secondary antibodies. Cells were imaged on an Echo Revolve microscope and quantified using ImageJ.

### (Supplemental)#Whole blood flow cytometry

Whole blood was obtained by cardiac puncture of *Cdh5-CreERT2; TdTomato^fl/fl^* animals at 12 weeks and 7 days old. 500 uL of whole blood was collected into tubes containing 40 uL of 0.5M EDTA (for P7, 5 animals were pooled to obtain enough blood), then spun down at 500g for 5 min. After removing supernatant, the pellet was resuspended in 9X volume of ammonium chloride solution (StemCell Technologies) and incubated on ice for 10 minutes for red blood cell lysis. As a positive control, a single cell suspension was made from 5 adult *Cdh5-CreERT2; TdTomato^fl/fl^* aortas by incubating in digestion solution (Liberase TH 70ug/mL, Sigma-Aldrich; DNase I, Qiagen; 10mM HEPES, Corning) at 37°C for 20 min before neutralizing with 10% FBS in DMEM. After pelleting, cells were then washed twice with 1X PBS before staining with CD45-FITC (BD) for 30 minutes on ice. Cells were then fixed in 4% PFA for 15 minutes before resuspending in FACS Buffer before analysis on a BD FACSymphony™ A5 Cell Analyzer.

## QUANTIFICATION AND STATISTICAL ANALYSIS

### (Supplemental)#Calculation of aorta volume

Aorta volume is calculated using the equation V = (C^2^ x h)/4π. V = volume, C = circumference of the aorta, h = length of the aorta.

### Preprocessing images for quantification

After optimization, images were cropped in FIJI along both the vertical and horizontal axes to 550 x 550 pixels to bring the number of objects to segment within a manageable working range.

### Nuclei segmentation

To extract endothelial nuclei and their spatial coordinates, we used deep-learning based cellpose (based on U-Net architecture) with the pre-trained model “nuclei”. If performance was suboptimal due to high background noise in individual images, we used “cyto2” instead. For hyperparameter optimization of the cellpose model parameters, we used the hyperparameter tuning techniques “manual search” and “grid search” to simultaneously decrease false positives and increase true positives; segmentation accuracy was validated by manually counting the percentage of true and false positives.

### Density analysis along the length and circumference axis

After processing through cellpose, stitched nuclei segmentations of whole aortas were divided into multiple sections of 500 x 200 pixels. The orientation was adjusted by averaging angles between the first and eighth intercostal regions. Using FIJI, the endothelium was segmented and intercostal vessel branch holes were subtracted for accuracy. Within each defined section of the aorta, we calculated the density by dividing the number of nuclei by the area of the section.

### Cell shape segmentation

To extract area, major axis, and minor axis at the individual cell level within the entire piece of tissue, we used segmentation based on PECAM signal of fixed and immunostained aortas. We leveraged Python library EPySeg ^58^, which provided accurate binary segmentation of endothelial cells. We transformed this output into an instance segmentation using the connected components function in python to enable analysis at the single-cell level. To ensure the reliability of our results, we excluded any segmentations that were less than 5 pixels away from the image border and verified that a minimum of 80% of the pixels of nucleus segmentation lay within the cell shape segmentation. To evaluate the quality of the segmentation, 20 randomly selected images were chosen and the quality of segmentation was verified manually. This method and associated quality control steps yielded 99% precision and 20% recall.

### Number of nuclei along length and circumference axes

On the basis of segmented nuclei, we created artificial strips along the length and circumference axes to estimate the number of cells in either direction (**Figure S2D-F**). Since the major axis represents cell length and the minor axis represents cell width, strips were aligned along the aorta’s length with a width equal to the median minor axis, and across its circumference with a width equal to the median major axis. To correct for alignment of the scanned tissue in the image, we determined the orientation angle by averaging the angles between the furthest nuclei of strips at both ends. The coordinates of the upper-left and lower-left corners of each strip were adjusted based on the angle and minor axis length. To get the coordinate of the tube on the right hand side, we use the formulas:

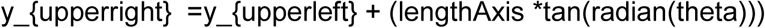

and

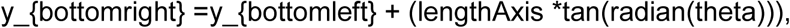

where tan() is the function to calculate the tangent value, radian() is the function to calculate the radian value, *theta* is the average angle of the orientation of the aorta, *y_{upperright }* is the upper coordinate of the tube on the right hand side, *y_{bottomright}* is the bottom coordinate of the tube on the right hand side, *y_{upperleft}* is the upper coordinate of the tube on the left hand side, and *y_{bottomleft}* is the bottom coordinate of the tube on the left hand side. Only nuclei with more than 75% within the strips were counted to avoid overestimation. Strips were iterated downwards in steps equal to half the major or minor axis until reaching the last nucleus on the image’s left side.

### Estimation of length and circumference

To estimate endothelium length from the first to the eighth intercostal, we multiplied the number of segmented nuclei along the length axis by the average major axis, then compared it to the measured length. For circumference, we multiplied the number of segmented nuclei along the circumferential axis by the average minor axis and compared it to the measured circumference.

### Endothelial nuclear signal (Ki67, EdU) extraction

To extract the percentage of nuclei that are Ki67 positive, we first merged previously generated nuclei segmentations with the Ki67 channel image to extract Ki67 intensities on the basis of nuclei coordinates. A background-based thresholding technique was used to distinguish real signal from noise, and we randomly selected 40 images across biological replicates and timepoints to validate the accuracy of this method. The same background-based thresholding technique was applied for EdU signal extraction as well. Using these methods, we achieved recall of 93% and 99% and precision of 98% and 99% for Ki67 and EdU, respectively. To remove prevalent Ki67 and EdU signal from smooth muscle cells, we filtered cell segmentation based on orientation, which successfully eliminated the majority of smooth muscle cell signals. Subsequent exclusion criteria based on nuclei size effectively filtered out all nuclei signals originating from smooth muscle cells.

### Mitotic spindle angle quantification

On NIS Elements software, a reference “zero angle” was defined as the line that connected two adjacent intercostal arteries. Mitotic spindle angle was determined by drawing a straight line between the two poles of the mitotic spindle (visualized by acetylated alpha-tubulin staining) and measuring the angle between the reference “zero angle” line.

### Clone size analysis of *Cdh5 Cre-ERT2; Rainbow^fl/+^* aortas

We decided to only analyze cells that had recombined to express mCherry or mOrange, since there was disproportionate overrepresentation of mCerulean cells. Each image was first split into one-channel images (nuclei, mOrange, and mCherry), and the mOrange or mCherry images underwent thresholding and watershed segmentation using python packages OpenCV and scikit-image. To define cell clusters, the resulting segmentations were connected to merge components that were touching each other. Simultaneously, we also performed nuclei segmentation as described previously. Nuclei segmentation was assigned to the cluster if >50% of the nuclei segmentation overlapped with the cluster signal. 5 biological replicates were randomly selected and evaluated; this method led to 96% precision and 95% recall.

### Regression analysis of *Cdh5 Cre-ERT2; Rainbow^fl/+^* aortas

To compare *Cdh5 Cre-ERT2; Rainbow^fl/+^* rainbow clone directions with tissue flow, we applied linear regression. For each clone, the regression line was calculated using the centroids of segmented nuclei within the clone.

### Nearest neighbor analysis

We conducted k-nearest neighbor analysis to explore spatial patterns of *Cdh5 Cre-ERT2; Rainbow^fl/+^* clone sizes. Centroids of each clone were identified, and distances to 1-5 nearest neighbors were calculated. We compared results with a random simulation and two biased simulations. For random simulation, nuclei were segmented, and n_cluster nuclei were randomly selected as centroids, with L2-distances measured over 1000 simulations. In biased simulations, the aorta was divided into three circumferential regions: clones appeared only in the middle for one simulation and in the upper and lower regions for the other. Random selection within the previously defined area and L2-distance calculations were repeated 1000 times. Distances were then normalized by the average distance from the random simulations.

### Gini coefficient analysis

We define *x*_*i*_, 1 ≤ *i* ≤ *n*, as the cluster size of cluster i, and n the number of identified clusters in one aorta. The Gini coefficient *G* is therefore calculated using the equation

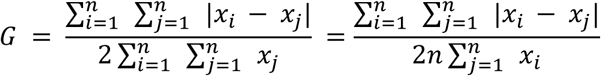

### Analysis of *Cdh5(cre); Rainbow^fl/+^* aortas with Ki67 immunofluorescence

mCherry and mOrange channels were segmented as previously described. Due to signal noise, we manually annotated and obtained coordinates of Ki67+ cells using FIJI software’s multi-point tool. For each segmented mCherry or mOrange cluster, we then determined the number of Ki67+ coordinates found within the cluster space. Without a nuclei immuno-staining (due to saturation of laser channels), Ki67+ cells required manual annotation in Fiji to obtain coordinates. To estimate the cluster sizes, we mapped rainbow signal area to cluster size based on previous inducible rainbow and nuclei segmentation data, adjusting for age dependencies over specified time intervals. We used time intervals of 0-5 and 0-10 to create a mapping for the interval 0-7. Similarly, we utilized time intervals of 5-30 and 5-60 to create a mapping for the interval 1-19. We excluded the time points of 21-30 from this analysis as the observed cluster sizes were significantly smaller than those visually observed in the interval 1-19.

### Distance analysis

Ki67-active cells were analyzed to determine their distribution across clusters grouped by cluster size. The experimental distribution was compared to simulations of three distinct scenarios: a Markovian division model, where each cell had an equal probability of being proliferative; a Markovian cluster model, where each cluster had an equal probability of containing a proliferative cell; and an exponential model, where the probability having a proliferative cell increased exponentially by cluster size. To quantify the similarity between experimental data and simulations, the distance was measured using three statistical metrics: Jensen-Shannon divergence, Wasserstein distance, and Kullback-Leibler divergence. For each scenario, distances were computed between the experimental data and 1,000 simulations. Additionally, pairwise distances were calculated between the 1,000 simulations within each scenario. Since these pairwise comparisons produced more than 1,000 distances, we sampled 1,000 distances as a negative control and compared these to the experimental data. For the first wave, clusters with sizes appearing fewer than ten times were filtered out before calculating the Jensen-Shannon divergence to ensure robust analysis.

### Generation of EdU lineage trees

EdU-active cells were identified as previously described for endothelial nuclei signal extraction. To establish “lineage” of an EdU cluster, daughter cells were required to share borders as confirmed by PECAM segmentation. Division trees were constructed based on EdU intensity and the cells’ position within the cluster, based on the assumption that limited movement occurs post-division.

### Mathematical modeling of aorta growth and cell loss

We defined two waves of cell divisions (*k*) that drive cellular proliferation and tissue growth (*k* ε {1, 2}). The first wave begins at postnatal day 3 and the second wave starts *T*_0_ days after the beginning of the first wave. Each wave is characterized by *C*_*k*_ rounds of division. The duration of cell cycle comes from an exponential distribution with a mean *T*_*k*_. In each division, *f*_*k*_ fraction of existing cells goes through division. We take into account cell loss by an effective rate of *P*_*k*_, which ensures that on an average, every division results into 2 − *P*_*k*_ cells. We calculate the total number of cells at time *t* after beginning of wave *k* as,

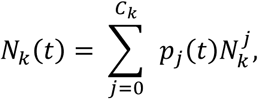

where, *p*_*j*_ is the probability of completion of *j* cell cycles calculated as,

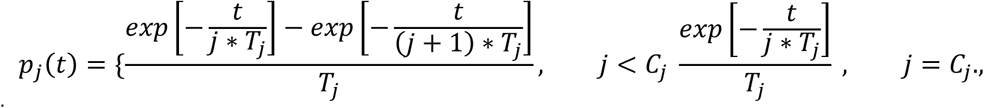

and 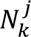 is the total number of cells in wave *k*, if there were definitely *j* divisions calculated as,

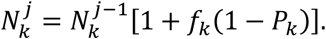

We set 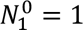 for comparison with the normalized experimental data and 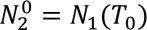.

We obtained the total number of cells going through active divisions as revealed by imaging experiments and quantification of Ki67+ nuclei.

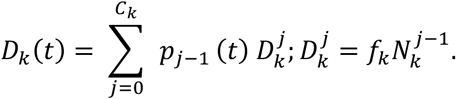

Finally, we calculate from our model, the number of clones of cells at time 6 which were already dividing at time 0. We find the probability of finding clones consisting of H cells at time 6 as,

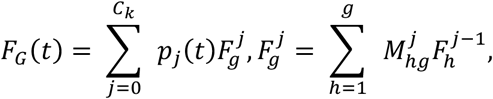

where, 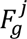 is the clone size distribution if exactly *j* divisions happen, and 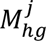 is the transition probability of clone size of ℎ resulting in clone size H in *j*^th^ division cycle calculated as,

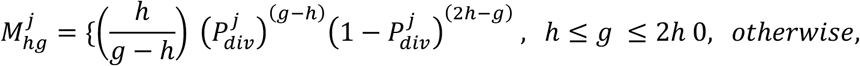

where, 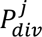 is the probability of a successful division. 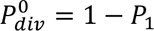, as all the marked cells are definitely dividing in the first cycle, whereas 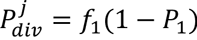 for *j* > 1. We use the parameters from the first wave for this calculation as the experimental data is within 3 days of marking. Altogether, this model has 9 free parameters (i.e. 2_’_, 4_%_, 3_%_, 2_%_, 5_%_, 4_+_, 3_+_, 2_+_, 5_+_). As this is a high-dimensional optimization problem, we use a Markov Chain Monte Carlo algorithm ^21^ that minimizes the difference between experimental observation and their equivalent from the model normalized by the experimental standard deviation ^38^. The initial guesses are taken from an uniform distribution in the range reported in **Table S3**.

To calculate cell cycle duration without cell loss (Figure 8E), 1000 Markov Chain Monte Carlo simulations were performed with *P_k_* set to 0. The frequency for each cell cycle duration (in units of days) is shown. To calculate what percentage of cells will complete cell division within a timeframe according to the mathematical model, we use the formula 1-exp(-*t* /effective cell cycle duration). For example, when *P_k_* has a nonzero value, median cell cycle duration is 1.45 days; 15.83% of cells within the model will complete division within 6 hours. Conversely, when *P_k_* = 0 and median cell cycle duration is 2.92 days, only 8.2% of cells within the model will complete division within 6 hours.

## REAGENTS

| REAGENT | CATALOG # |
| --- | --- |
| **Antibodies** |  |
| Rabbit anti-ERG recombinant monoclonal antibody | ab92513 |
| Rabbit anti-ERG (SP06-04) recombinant monoclonal antibody | Thermo Fisher #MA5-32036 |
| Armenian Hamster anti-PECAM (2H8) monoclonal antibody | Developmental Studies Hybridoma Bank |
| Rabbit anti-Ki67 [SP6] | ab16667 |
| Rabbit Alexa Fluor® 647 Anti-Ki67 antibody [SP6] | ab281928 |
| Rabbit anti-acetylated alpha tubulin (Lys40) (D20G3) XP® | Cell Signaling Technology #5335 |
| Chicken anti-GFP polyclonal antibody | ab13970 |
| Rabbit Living Colors® DsRed polyclonal antibody | Takara Bio #632496 |
| Mouse Alexa Fluor® 647 HA-Tag (6E2) monoclonal antibody | Cell Signaling Technology #3444 |
| FITC Rat anti-Mouse CD45 | BD Biosciences #553080 |
| **Chemicals** |  |
| Tamoxifen | MP Biochemicals #MP215673891 |
| Tamoxifen | Sigma Aldrich #T5648 |
| 5-Ethynyl-2'-deoxyuridine | Med Chem Express #50-225-9229 |
| Sunflower seed oil | Sigma Aldrich #S5007 |
| Castor oil, ethoxylated | Sigma Aldrich #C5135 |
| Liberase TH | Roche #5401135001 |
| DNase I | Qiagen #79254 |
| HEPES | Sigma Aldrich #H0887 |
| Trypsin | Gibco #15400054 |
| **Commercial Kits** |  |
| Vasculpaint Ex Vivo Contrast Agent | Medlumine Cat #MDL-121 |
| Click-iT™ EdU Cell Proliferation Kit for Imaging, Alexa Fluor™ 594 dye | Invitrogen #C10339 |
| **Mouse Strains** |  |
| Cdh5(PAC)-CreERT2 (inducible VE-cadherin-Cre) | Taconic Biosciences #3848982 |
| BVEN-Cre (constitutive VE-cadherin-Cre) | Jackson Laboratories #006137 |
| Osx1-GFP; Cre | Jackson Laboratories #006361 |
| Rainbow fl/fl | Laboratory of Dr. Irving Weissman |
| TdTomato fl/fl | Jackson Laboratories #007914 |
| iMb2-Control-Mosaic; iChr2-Control-Mosaic | Jackson Laboratories #031298 |
| Fgfr1 fl/fl | Laboratory of Dr. Juha Partenen |
| Fgfr2 fl/fl | Jackson Laboratories #007569 |
| PIP-FUCCI fl/fl | Laboratory of Dr. Victoria Bautch |
| C57BL/6J | Jackson Laboratories #000664 |

## SUPPLEMENTAL FIGURE LEGENDS

**Supplemental Figure 1.**

(A) Measured length (mm) of descending aorta between the 1st - 8th intercostal vessels, with males (blue) and females (yellow) indicated. Inset shows the rates of growth of aorta length from postnatal day 3 to 7 (P3-P7) and from P7 to P14. Each dot represents a biological replicate, with the mean and standard deviation shown. N = 5-7 animals per age.

(B) Vertebral length (black), aorta circumference (orange), and aorta length (pink) normalized to minimum values.

(C) Aorta volume (blue) and animal weight (pink) normalized to minimum values.

(D) Representative microcomputed tomography reconstructions of backbones from wildtype (*Fgfr1^fl/fl^* and *Fgfr2^fl/fl^*) and mutant (*Osx-cre; Fgfr1^fl/fl^ Fgfr2^fl/fl^*) animals. The arrows indicate the position of C1 and L1 vertebrates. Scale bar = 2mm.

**Supplemental Figure 2.**

(A) Additional examples of stitched whole mount confocal images of aortas from each age. ERG (red) and PECAM (white). Scale bar = 500 µm.

(B) Representative cross-section confocal images of aortas from each age to show the height of cells. ERG (red) and PECAM (white). Scale bar = 50 µm.

(C) Schematic of quality check of cell shape segmentation using cell shape segmentation, nucleus segmentation, and criteria for inclusion.

(D) Schematic showing the method of determining the angle of blood flow θ. Nuclei are visualized by ERG immuno-staining (white).

(E) Schematic showing the method of counting the number of cells along the length axis. Nuclei are visualized by ERG immuno-staining (white).

(F) Schematic showing the method of counting the number of cells along the circumference axis. Nuclei are visualized by ERG immuno-staining (white).

**Supplemental Figure 3.**

(A) Schematic showing segmentation method for Ki67 or EdU nuclear signal, and the criteria for inclusion. Ki67 or EdU signal was only recorded if signal_inside >threshold_inside, and signal_inside / signal_outside > threshold_fraction.

(B) Ki67 and ERG *en face* whole mount immunostaining from P18, P19, P20, P21, P22, P23 animals. Each image is a biological replicate. Scale bar = 100 µm.

(C) Example of how mitotic spindle angle was measured in NIS Elements. The blue dashed line indicates the direction of flow. Scale bar = 20 µm.

(D) Additional representative images showing active mitotic spindles in aorta endothelium. Scale bar = 10 µm.

**Supplemental Figure 4.**

(A) Frequency and expression level of cells expressing Tagln, CD3e, Dcn, Pecam1, or Prox1. The red dashed line indicates the cutoff where cells below cutoff were given a score of 0, and celow above the cutoff were given a score of 1.

(B) UMAP of overall cell doublet scoring, which is the combined score from Tagln, CD3e, Dcn, Pecam1, or Prox1.

(C) UMAP of cells that were identified as singlets or non-singlets. Singlets were defined as cells that had a score < 2 and were kept for further analysis. Non-singlets were identified as cells that had a score > 2 and were removed from further analysis

(D) Elbow plot indicating the standard deviation of principal components (PC). The cutoff is shown at 30.

(E) Percentage of proliferating endothelial cells from single cell RNA sequencing data.

(F) Violin plots of Acetyl CoA score from P5, P10, P20, and P30 endothelial cells. The score of the 90^th^ percentile is indicated in numbers and with a dashed line.

(G) UMAPs showing expression of Acetyl CoA genes in P5, P10, P20, and P30 endothelial cells.

(H) UMAPs showing expression of fatty acid genes in P5, P10, P20, and P30 endothelial cells.

(I) UMAPs showing expression of extracellular matrix genes in P5, P10, P20, and P30 endothelial cells.

(J) Heat map showing top differentially expressed genes between P5, P10, P20, and P30 endothelial cells.

**Supplemental Figure 5.**

(A) Representative images from constitutive lineage tracing animals aged P10, P30, and P90. Each image is a biological replicate. Scale bar = 100 µm.

(B) (Top) Dosing scheme for tamoxifen. (Bottom) Representative images from inducible lineage tracing animals after having received 1 dose, 2 doses, or 4 doses of tamoxifen. Each image is a biological replicate.

(C) Total tagged clones in the red (mCherry) and orange (mOrange) channel from inducible lineage tracing animals, across induction and harvest time windows. The mean and standard deviation are indicated. N = 4-6 animals per age.

(D) The fraction of dividing clones in the red (mCherry) and orange (mOrange) channel from inducible lineage tracing animals, across induction and harvest time windows. The mean and standard deviation are indicated. N = 4-6 animals per age.

(E) Average clone size in the red (mCherry) and orange (mOrange) channel from inducible lineage tracing animals, across induction and harvest time windows. The mean and standard deviation are indicated. N = 4-6 animals per age.

(F) Frequency of clone size in the red (mCherry) and orange (mOrange) channel from inducible lineage tracing animals, in P5-P30 and P30-P60 time windows. N = 4-6 animals per age.

(G) Schematic of iMb2-Mosaic; iChr2-Mosaic (expression driven by Cdh5 Cre) lineage tracing construct.

(H) Possible clone combinations from iMb2-Mosaic; iChr2-Mosaic lineage tracing.

(I) Representative images of iMb2-Mosaic; iChr2-Mosaic clones within the first wave (P3-P10) and second wave (P20-P27). Representative clones are outlined using dashed lines.

(J) Representative images of en face whole mount aorta from inducible lineage tracing animals, across time windows studied. Each image is a biological replicate. Scale bar = 200 µm.

(K) Clone distributions from each biological replicate within each time window, with singlets included.

(L) Clone distributions from each biological replicate within each time window, with singlets excluded.

(M) Normalized distance to the nearest neighbor of each red colored clone from each time window is represented, with each dot corresponding to a cluster and each color indicating a biological replicate. The mean and standard deviation is shown.

(N) Normalized distance to the nearest neighbor is shown for red and orange-colored clones from each time window, with each dot representing a cluster. The number of biological replicates are shown in the data. The mean and standard deviation is shown.

(O) Normalized distance of the first to the fifth nearest neighbor of each red colored clone from each time window is represented, with each dot corresponding to a cluster. The number of biological replicates are shown in the data. The mean and standard deviation is shown.

(P) Average proportion of cluster sizes including singlets, found across all induction and harvest time windows.

(Q) Average clone size including singlets from each time window; the color of the bar represents the maximum clone size found within that time window. N = 5-8 animals per time point.

(R) Average coefficient of variation of clone size from each time window; the color of the bar represents the maximum clone size found within that time window. N = 5-8 animals per time point.

**Supplemental Figure 6.**

(A) Gini coefficient for average cluster size, percentage of proliferating tagged cells, and percentage of proliferating tagged clones. Blue dots indicate time windows found within the first wave, gray dots indicate time windows found within the second wave, and orange dots indicate time windows found within both. N = 5-8 animals per age.

(B) Hoover index, as above.

(C) Shannon Log2 equilibrium, as above.

(D) Shannon LogN equilibrium, as above.

(E) Theil index, as above.

(F) KL divergence of experimental and simulated data for scenario 1, scenario 2, and scenario 3 for the first wave.

(G) Wasserstein distance of experimental and simulated data for scenario 1, scenario 2, and scenario 3 for the first wave.

(H) Jensen Shannon distance of experimental and simulated data for scenario 1, scenario 2, and scenario 3 for the first and second wave.

(I) KL divergence of experimental and simulated data for scenario 1, scenario 2, and scenario 3 for the first and second wave.

(J) Wasserstein distance of experimental and simulated data for scenario 1, scenario 2, and scenario 3 for the first and second wave.

**Supplemental Figure 7.**

(A) (Top) Experimental design for measuring EdU activity after IP injection into mice. (Bottom) Representative images of fibroblasts treated with fresh EdU, or serum harvested from mice 20 minutes, 60 minutes, or 180 minutes post-EdU IP injection. Red arrows point to EdU+ cells and white asterisks indicate Ki67+ cells.

(B) Percentage of Ki67+ (green) and EdU+ (red) cells from fibroblasts treated with fresh EdU, or serum harvested from mice 20 minutes, 60 minutes, or 180 minutes post-EdU injection.

**Supplemental Figure 8.**

(A) Experimental data (solid) and model simulated data (dashed) of fraction of dividing cells over time. The mean and standard deviation are shown.

(B) Experimental data (solid) and model simulated data (dashed) of the ratio of clones to singlets over time. The mean and standard deviation are shown.

(C) - (E) 2D histograms showing the frequency of optimal parameters from 1000 iterations of MCMC simulation. The pair of parameters in the respective panels are (C) f_1_ (fraction of existing cells dividing in wave 1) and T_1_ (duration of cell cycle in wave 1, in days); (D) f_2_ (fraction of existing cells dividing in wave 2) and T_2_ (duration of cell cycle in wave 2, in days); and (E) P_1_ (probability of cell extrusion in wave 1) and P_2_ (probability of cell extrusion in wave 2).

(F) The proportion of clusters across time with 1, 2, 3, 4, or 5+ cells, after EdU injection in either the first (injected at P5) or second wave (injected at P19).

(G) The normalized distance between all clusters, between singletons and clusters, between cells within a cluster, and between cells that are 2 cells apart within a cluster, for clones observed 48 hours after EdU injection. The mean and standard deviation are shown.

(H) As above, for clones observed 72 hours after EdU injection. The mean and standard deviation are shown.

(I) Representative image showing an endothelial cell extruding above the monolayer. Scale bar = 50 µm.

(J) Cross sections of endothelial cell extruding above the monolayer. Scale bar = 25 µm.

(K) Additional example images of EdU singlets that are small in size. Singlets are outlined with dashed lines. Scale bar = 50 µm.

(L) Gating strategy for cells dissociated from 5 adult *Cdh5-Cre; tdTomato^fl/fl^* aortas.

(M) Gating strategy for cells collected from 500 uL whole blood from 1 adult *Cdh5-Cre; tdTomato^fl/fl^* animal.

(N) As above, an additional biological replicate.

(O) As above, an additional biological replicate.

(P) Gating strategy for cells collected from 500 uL pooled whole blood from 5 first wave (P7) *Cdh5-Cre; tdTomato^fl/fl^* animals.

**Supplemental Figure 9**

(A) MicroCT cross-sections and identification of vertebrae.

(B) 3D reconstructions of microCT with S1 vertebrae identified. The caudal position is towards the top of the image.

(C) 3D reconstructions of microCT with L6, L5, L4 vertebrae and their respective vertebral centrum’s cranial Z positions identified.

