## Supplemental Figures 1-9 for "Resolving the design principles that control postnatal vascular growth and scaling"

### Supplementary Figure 1

Click here to access/download;Supplemental figure;RelatedToMainFigure1.pdf

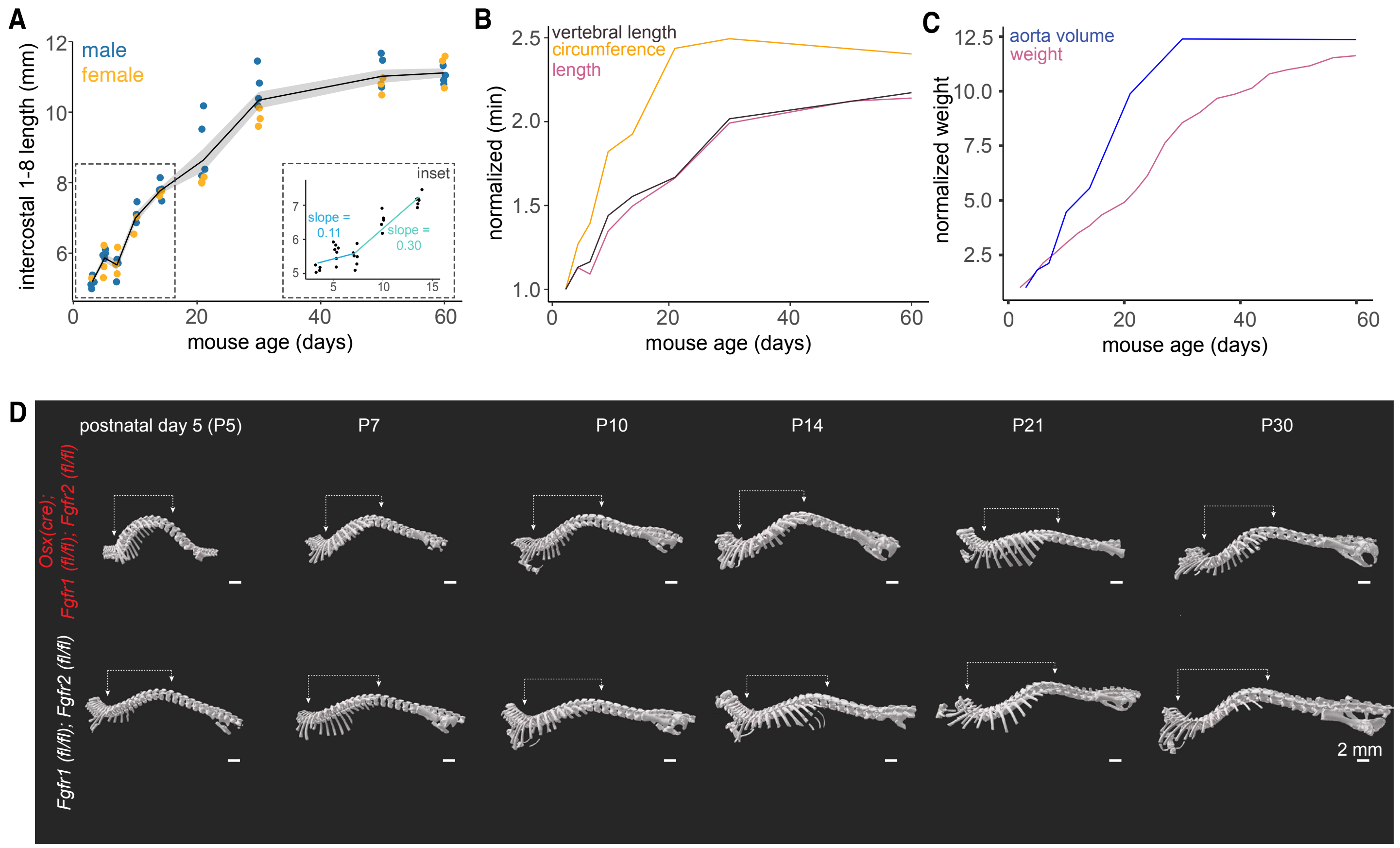

Click here to access/download;Supplemental figure;RelatedtoMainFigure2_1.pdf

#### Supplementary Figure 2

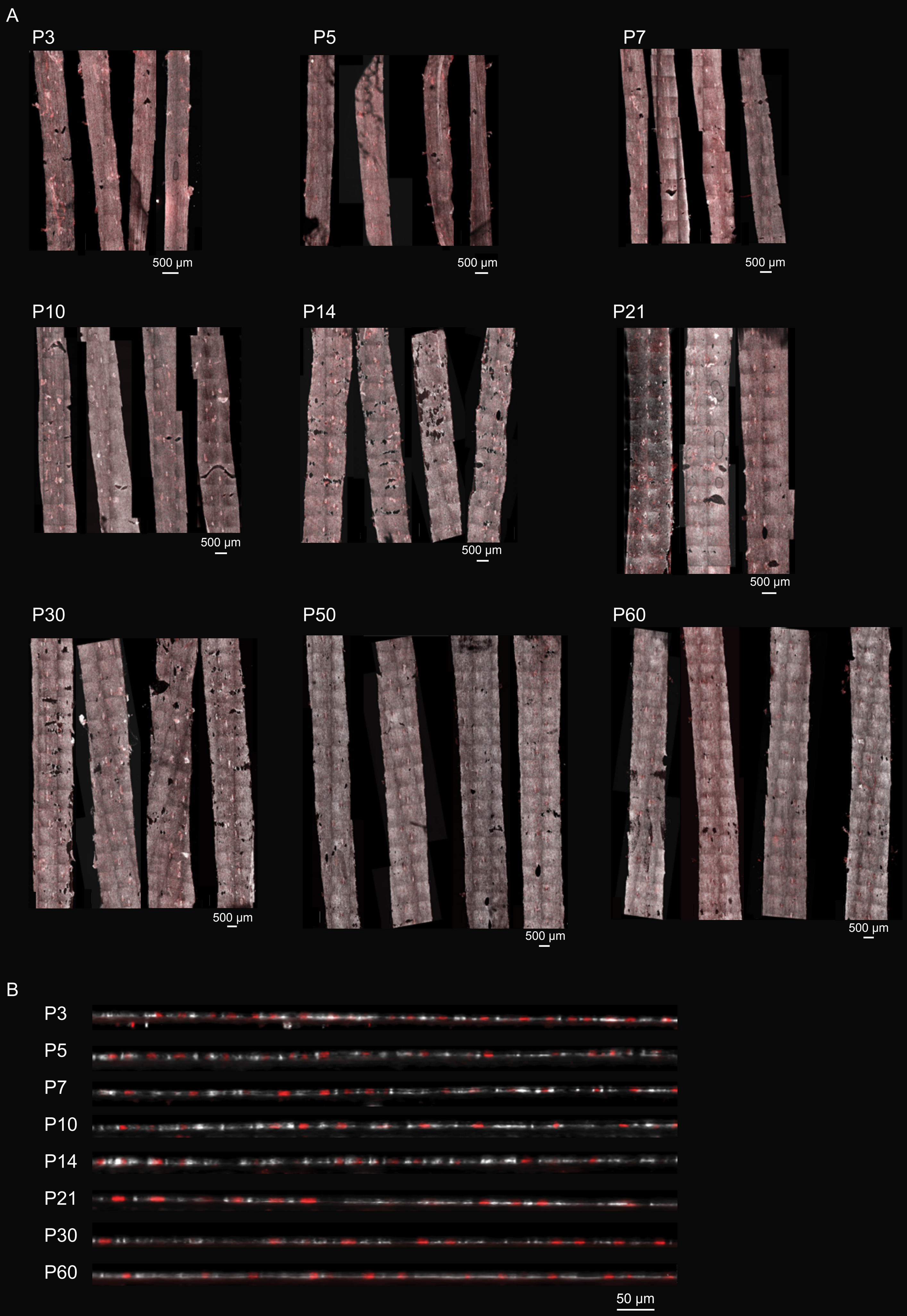

Supplemental Figure 2
Click here to access/download;Supplemental figure;RelatedtoMainFigure2_2.pdf

### Supplementary Figure 2

C

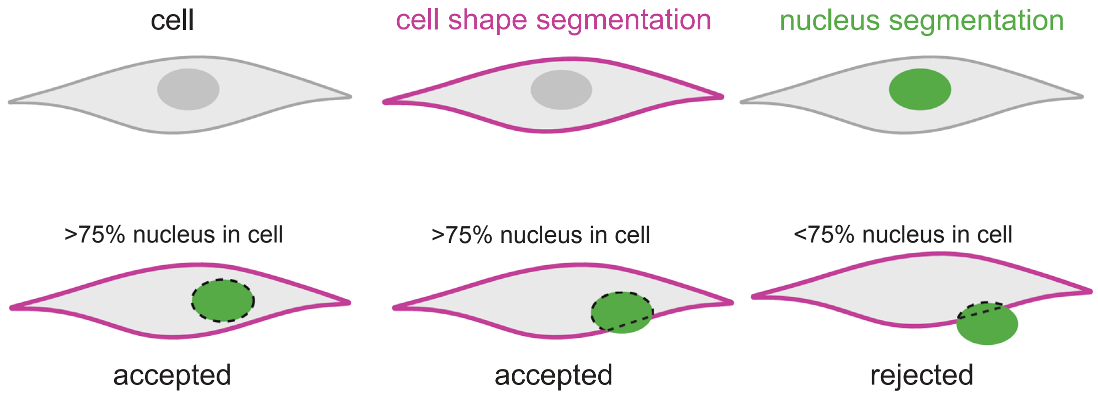


**D** Determining angle of blood flow

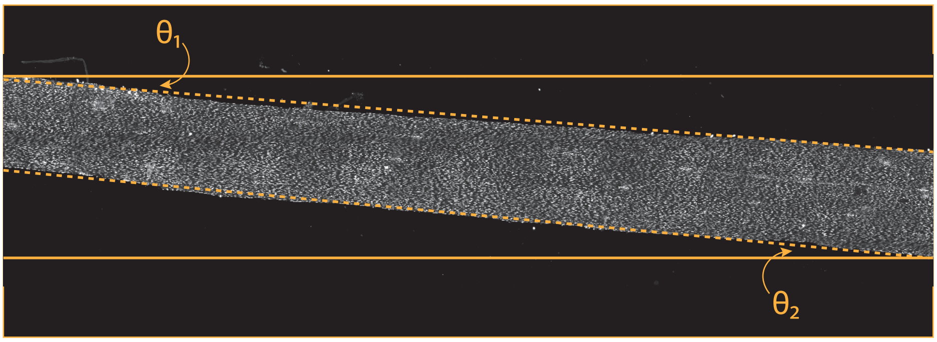


Angle of blood flow ($\theta$)

$\theta = (\theta_1 + \theta_2) / 2$

**E** Counting number of cells along length axis

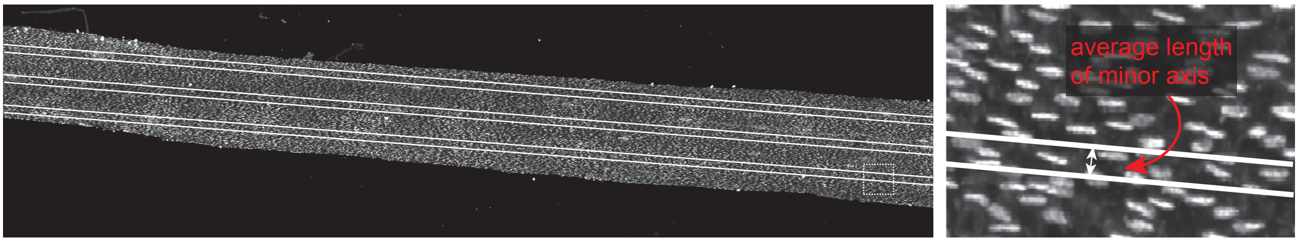


**F** Counting number of cells along circumference axis

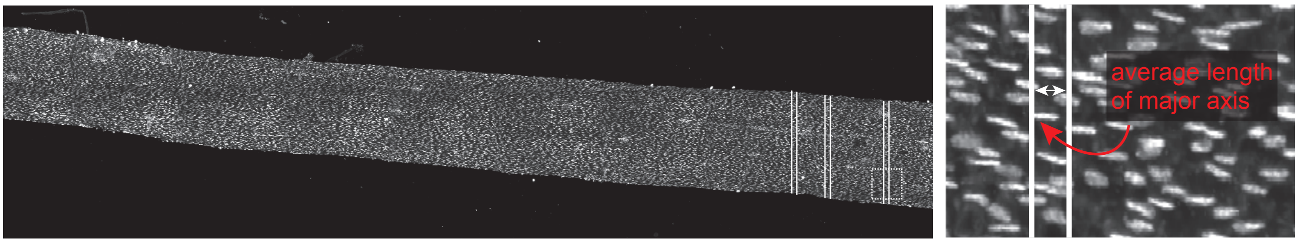

Supplemental Figure 3

Click here to access/download;Supplemental figure;RelatedtoMainFigure3.pdf

**Supplementary Figure 3**

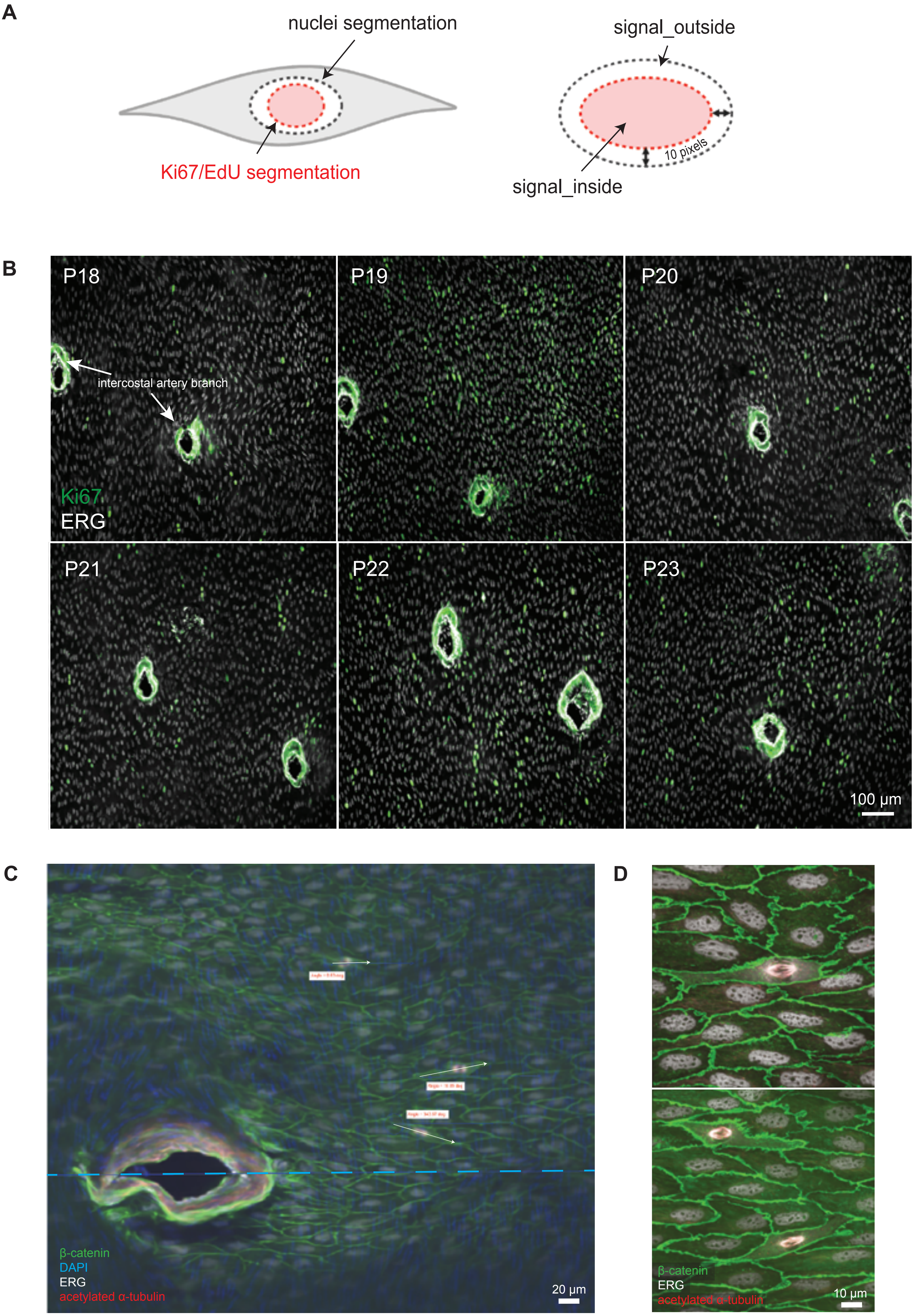

Supplemental Figure 4

Click here to access/download;Supplemental figure;RelatedtoMainFigure4.pdf

**Supplementary Figure 4**

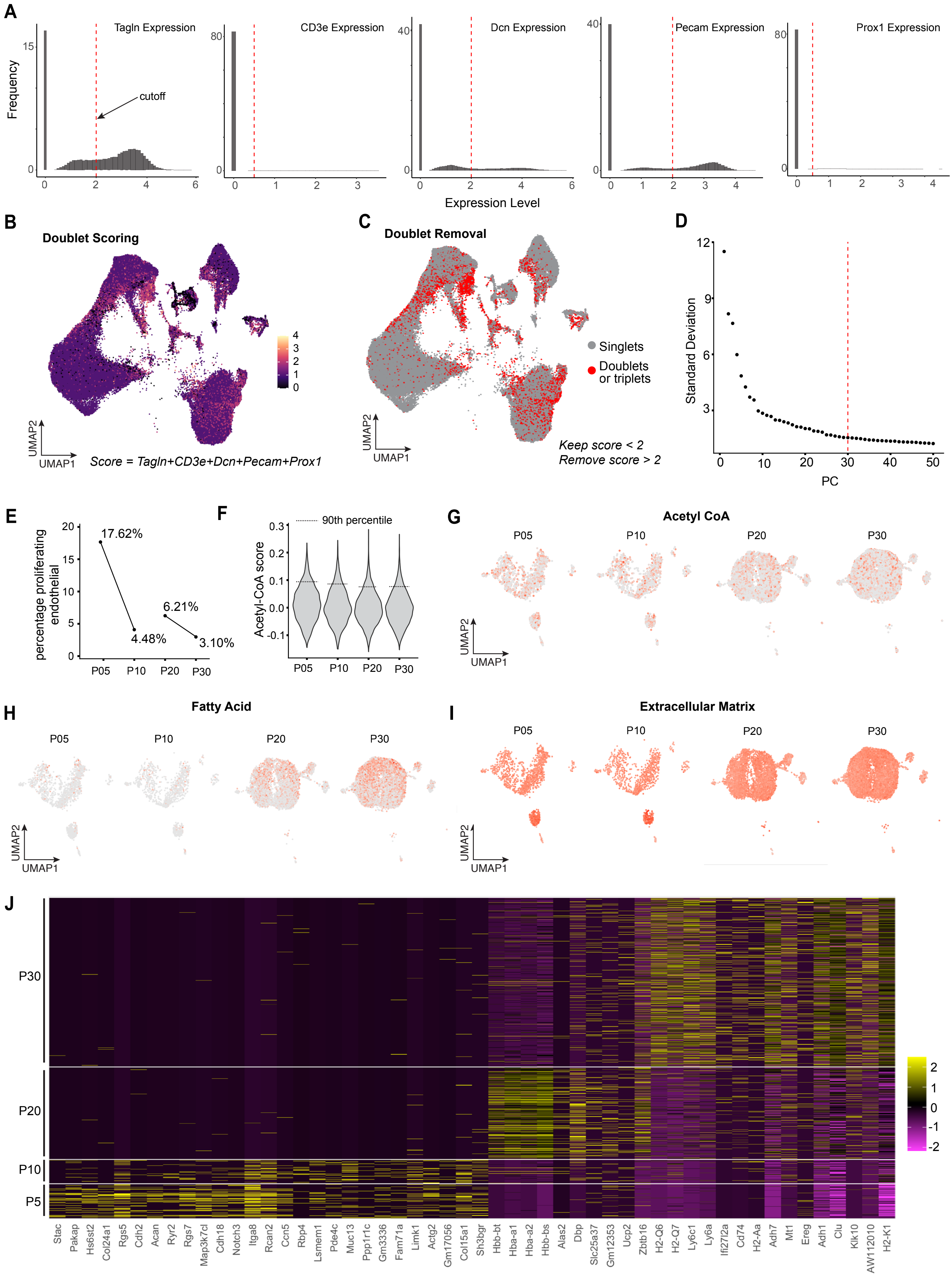

Supplemental Figure 5
Click here to access/download;Supplemental figure;RelatedtoMainFigure5_1.pdf

**Supplementary Figure 5**

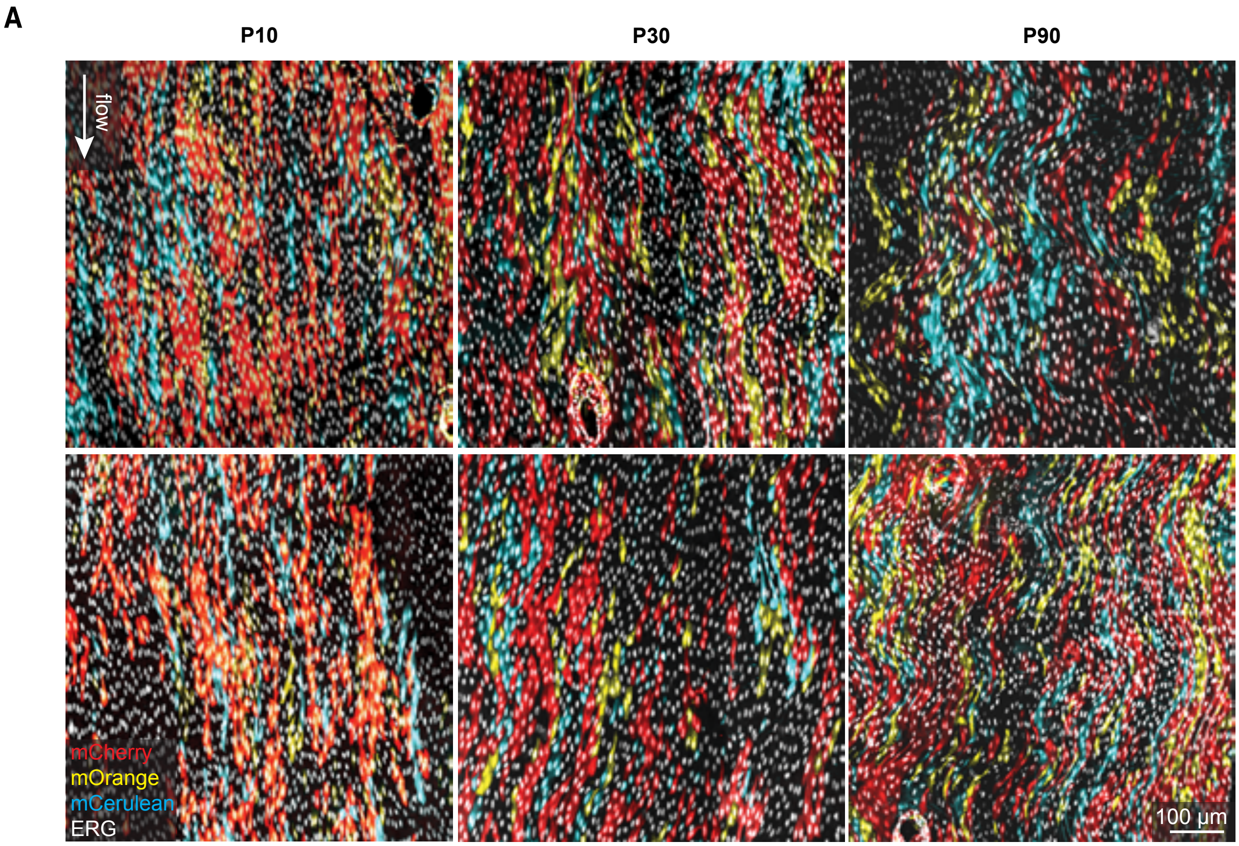


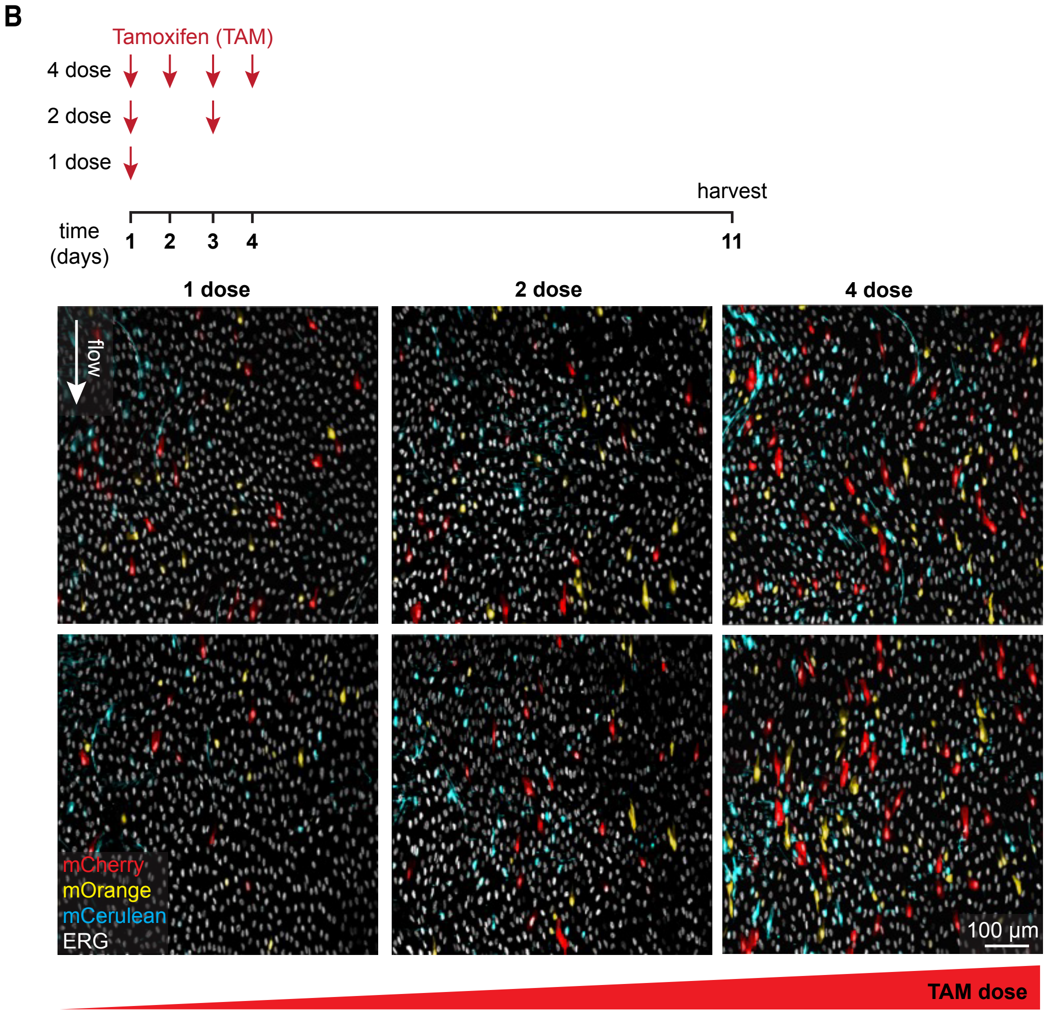

Supplemental Figure 5

Supplementary Figure 5

Click here to access/download;Supplemental figure;RelatedtoMainFigure5_2.pdf

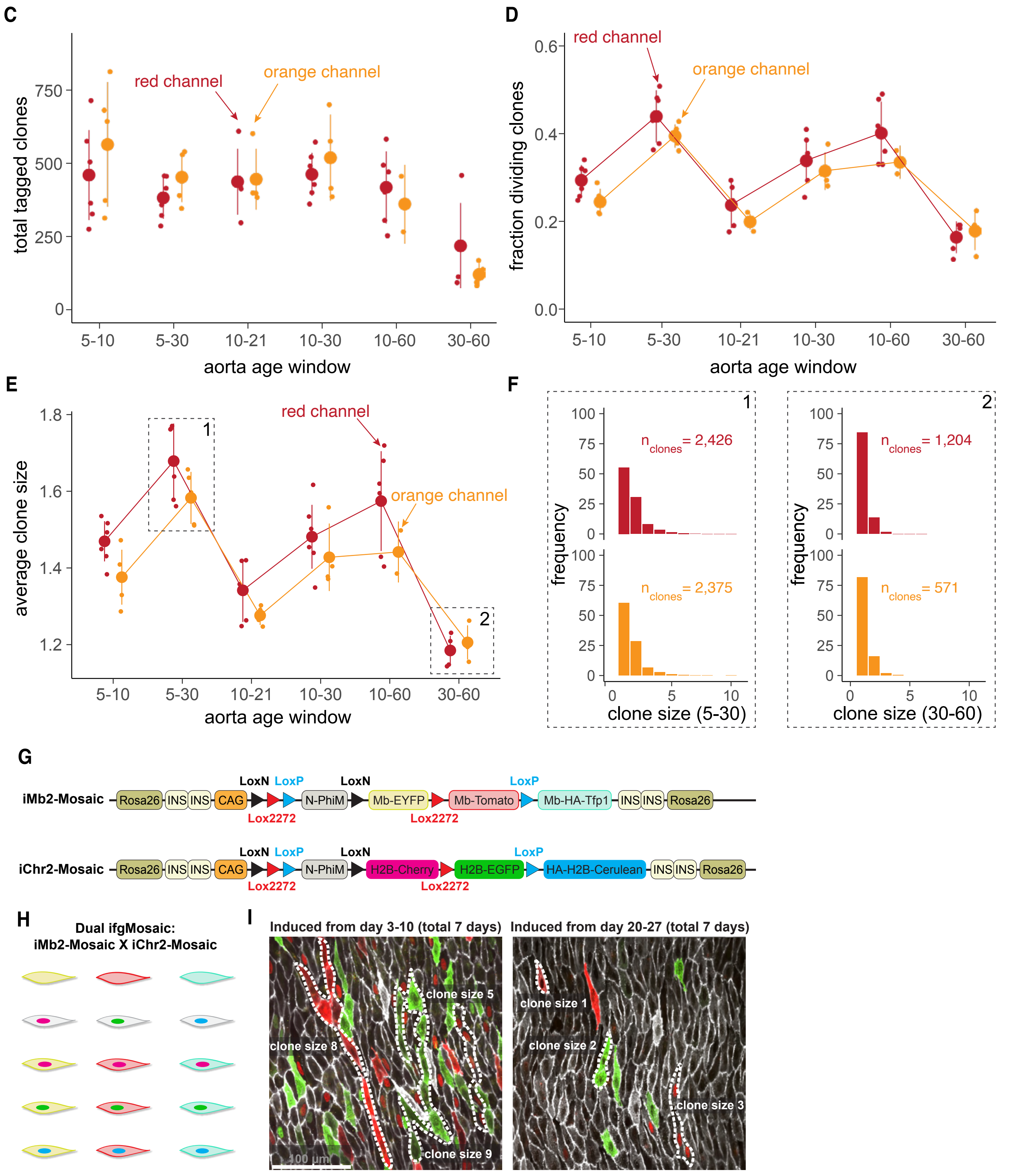

Click here to access/download;Supplemental figure;RelatedtoMainFigure5...pdf

Supplementary Figure 5

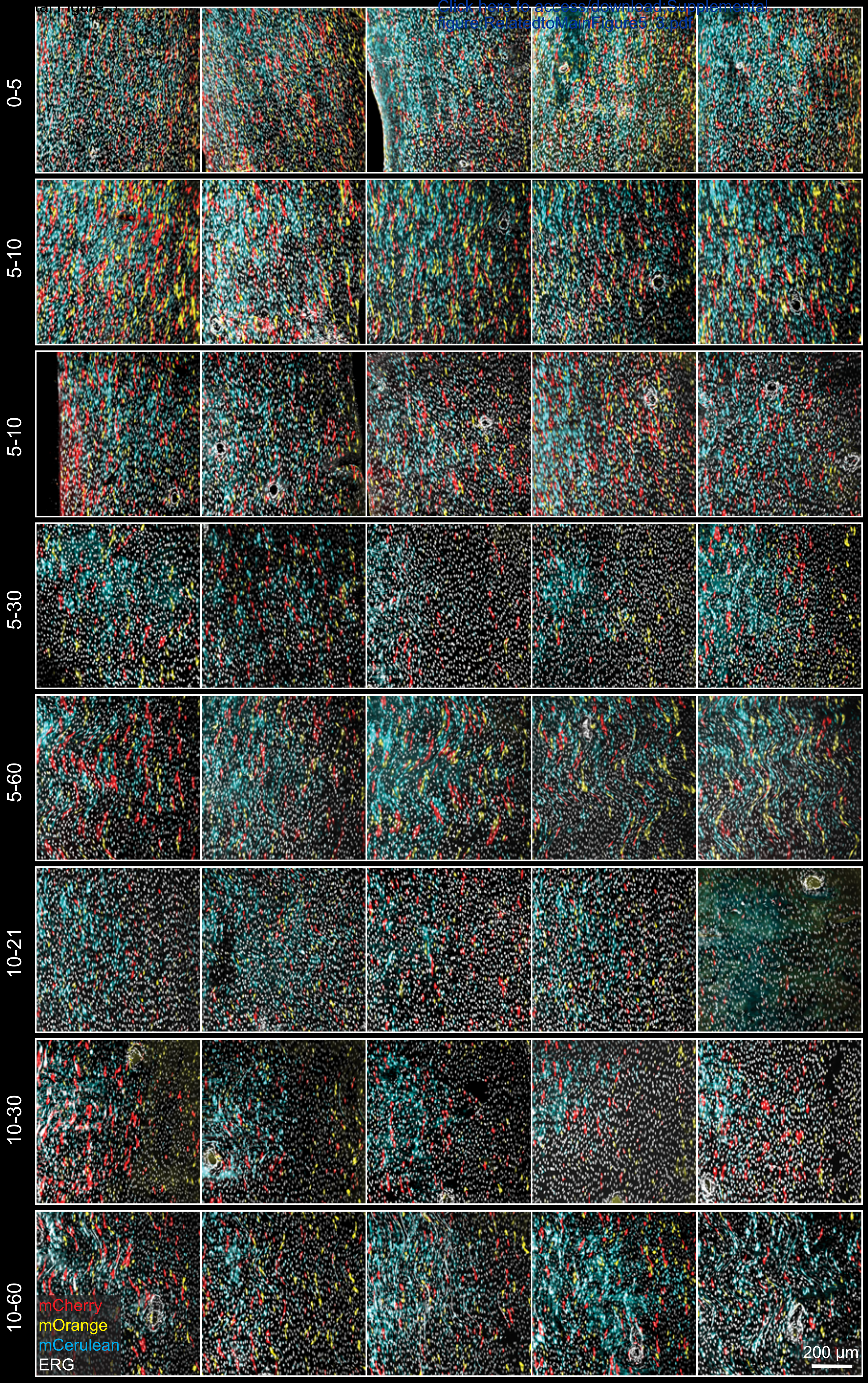

Supplemental Figure 5

### Supplementary Figure 5

Click here to access/download;Supplemental figure;RelatedtoMainFigure5_4.pdf

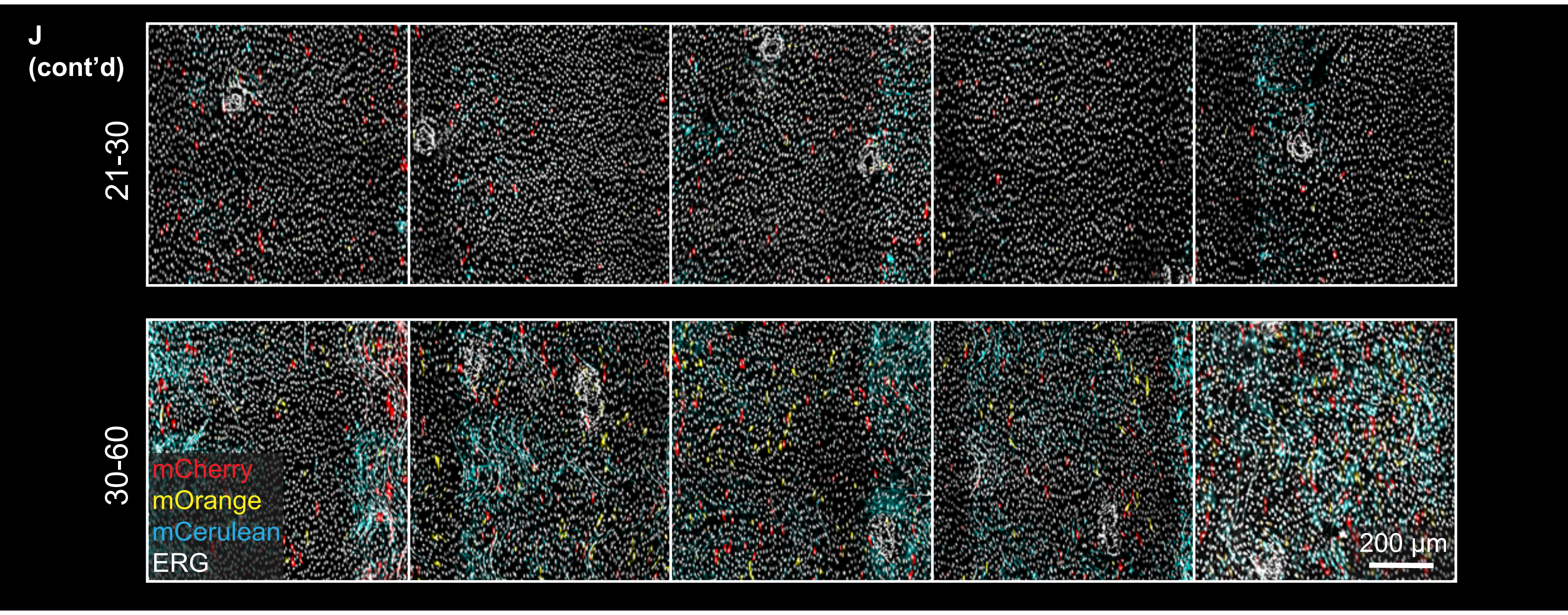


K

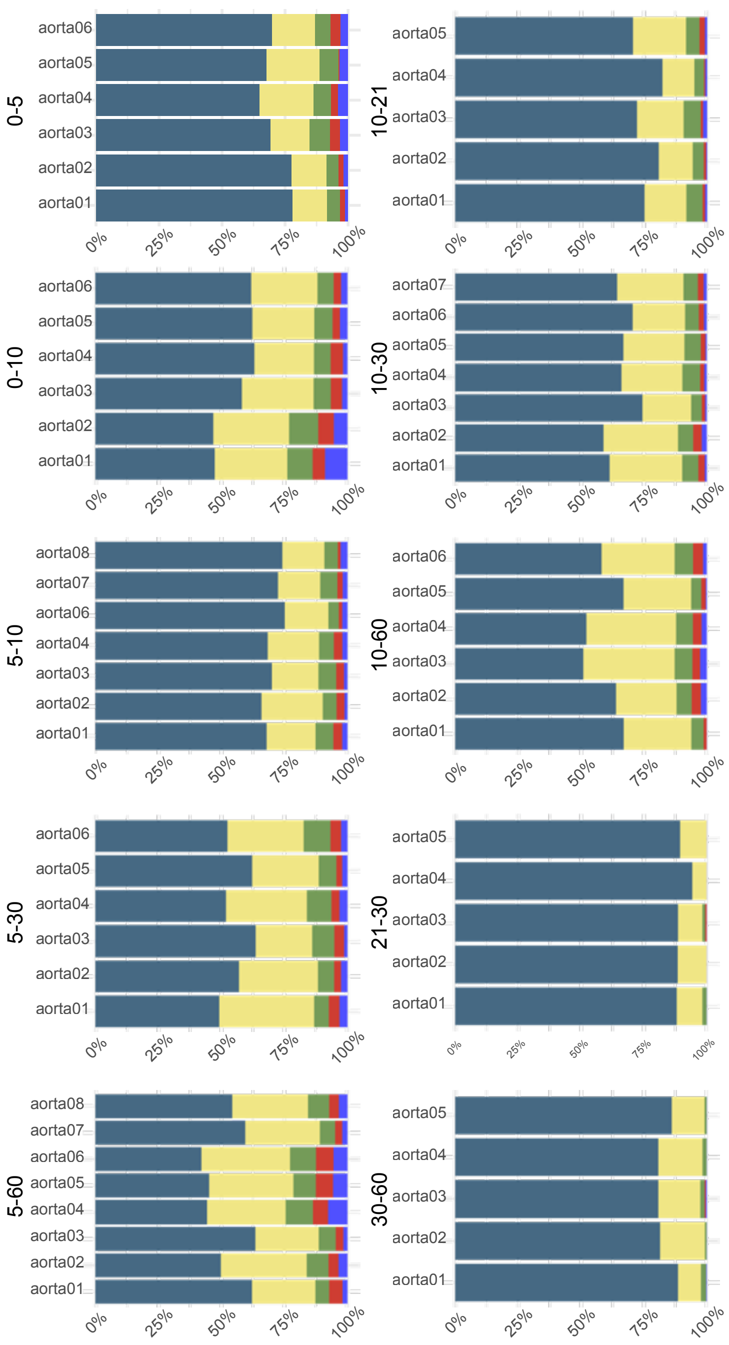


L

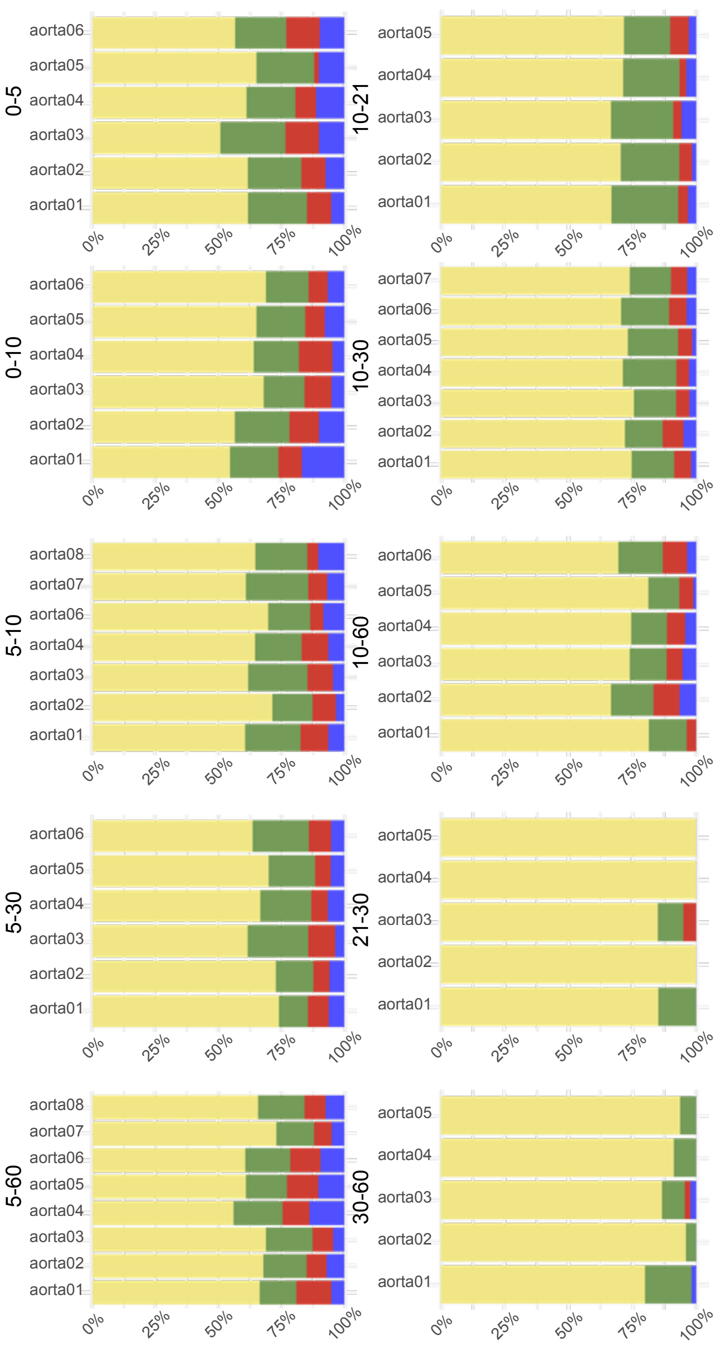

Supplemental Figure 5

### Supplementary Figure 5

Click here to access/download;Supplemental figure;RelatedtoMainFigure5_5.pdf

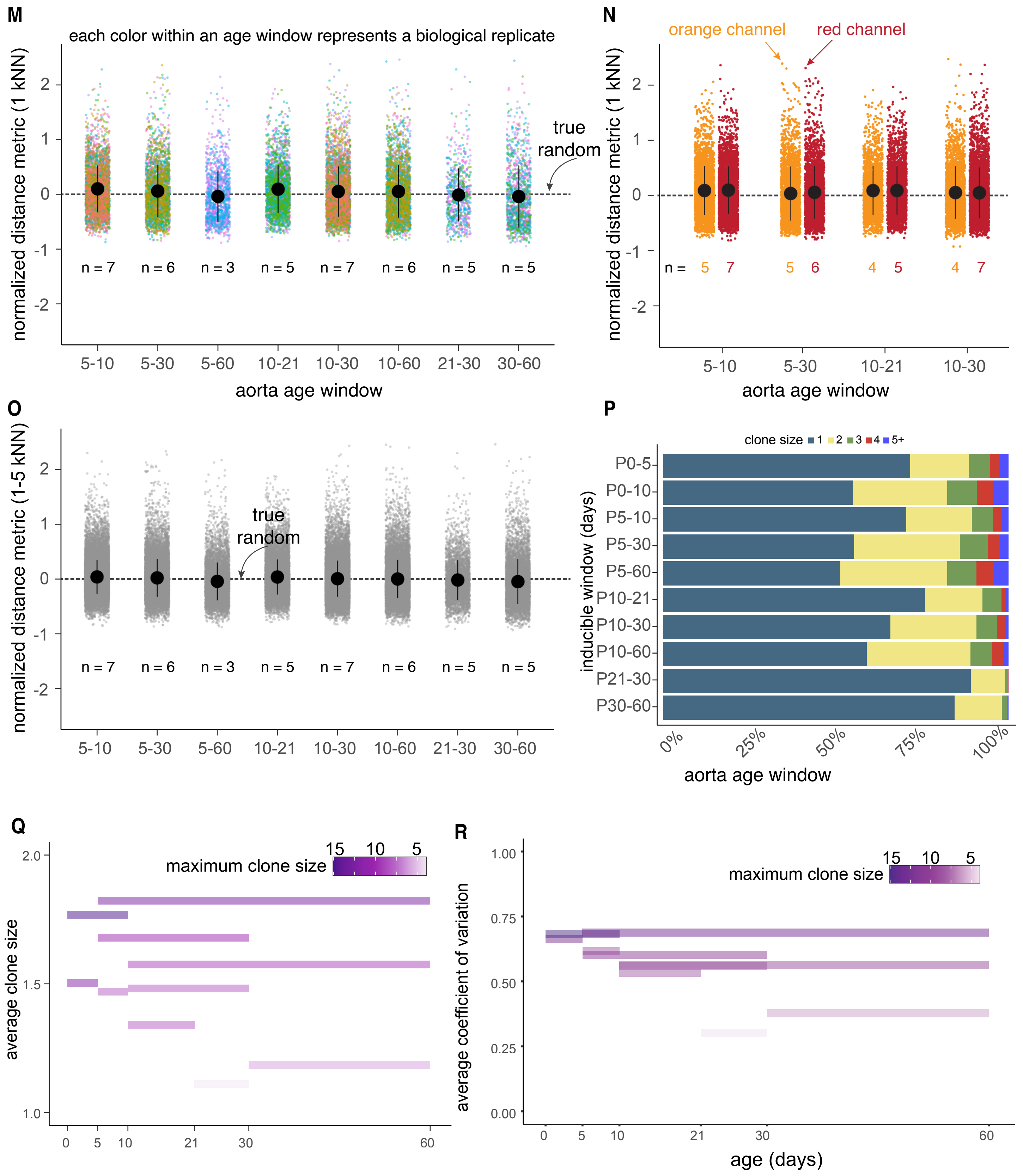

Supplemental Figure 6

### Supplementary Figure 6

Click here to access/download;Supplemental figure;RelatedtoMainFigure6_1.pdf

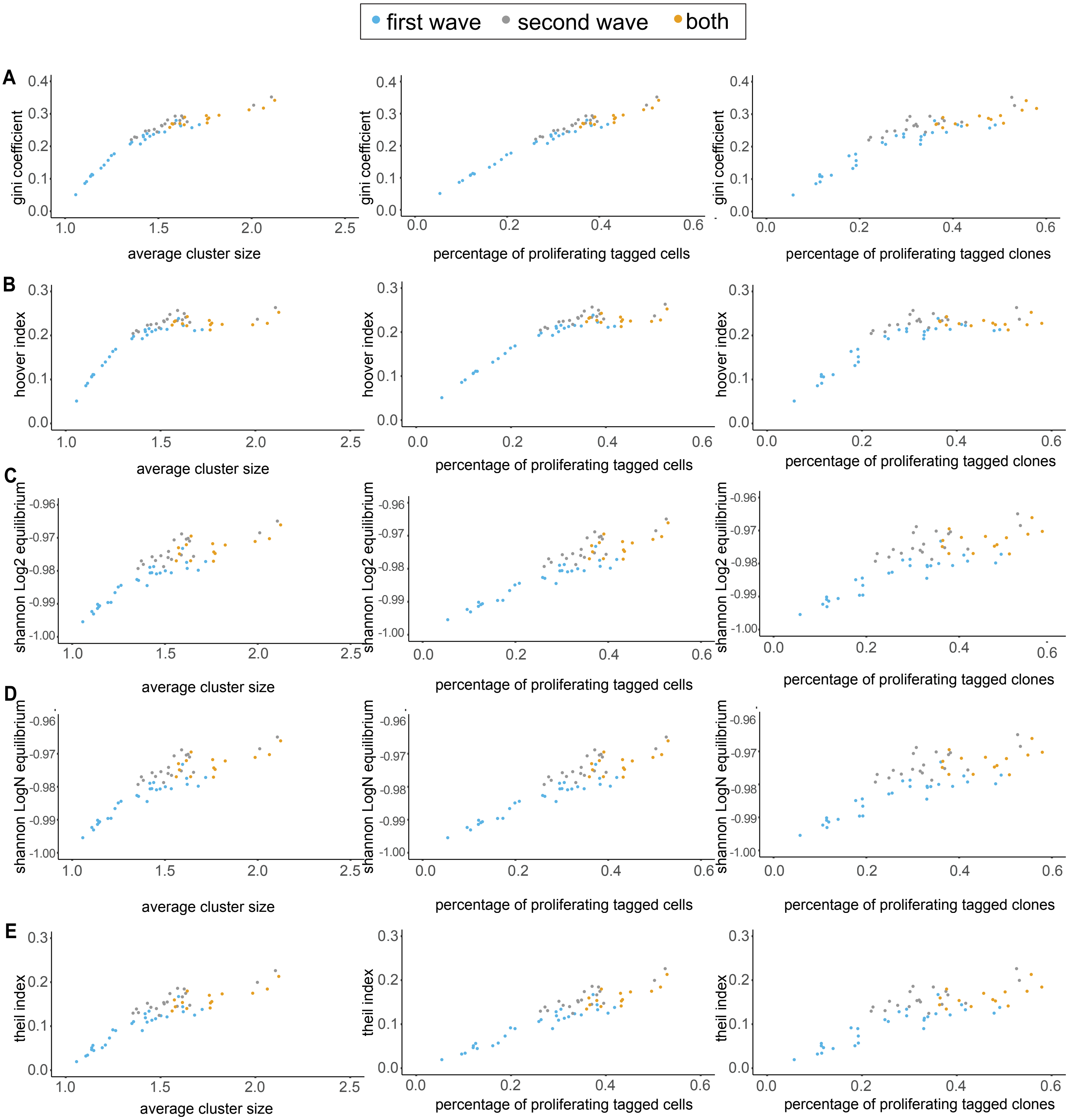

**Supplementary Figure 6**

Supplemental Figure 6

Click here to access/download;Supplemental figure;RelatedtoMainFigure6_2.pdf

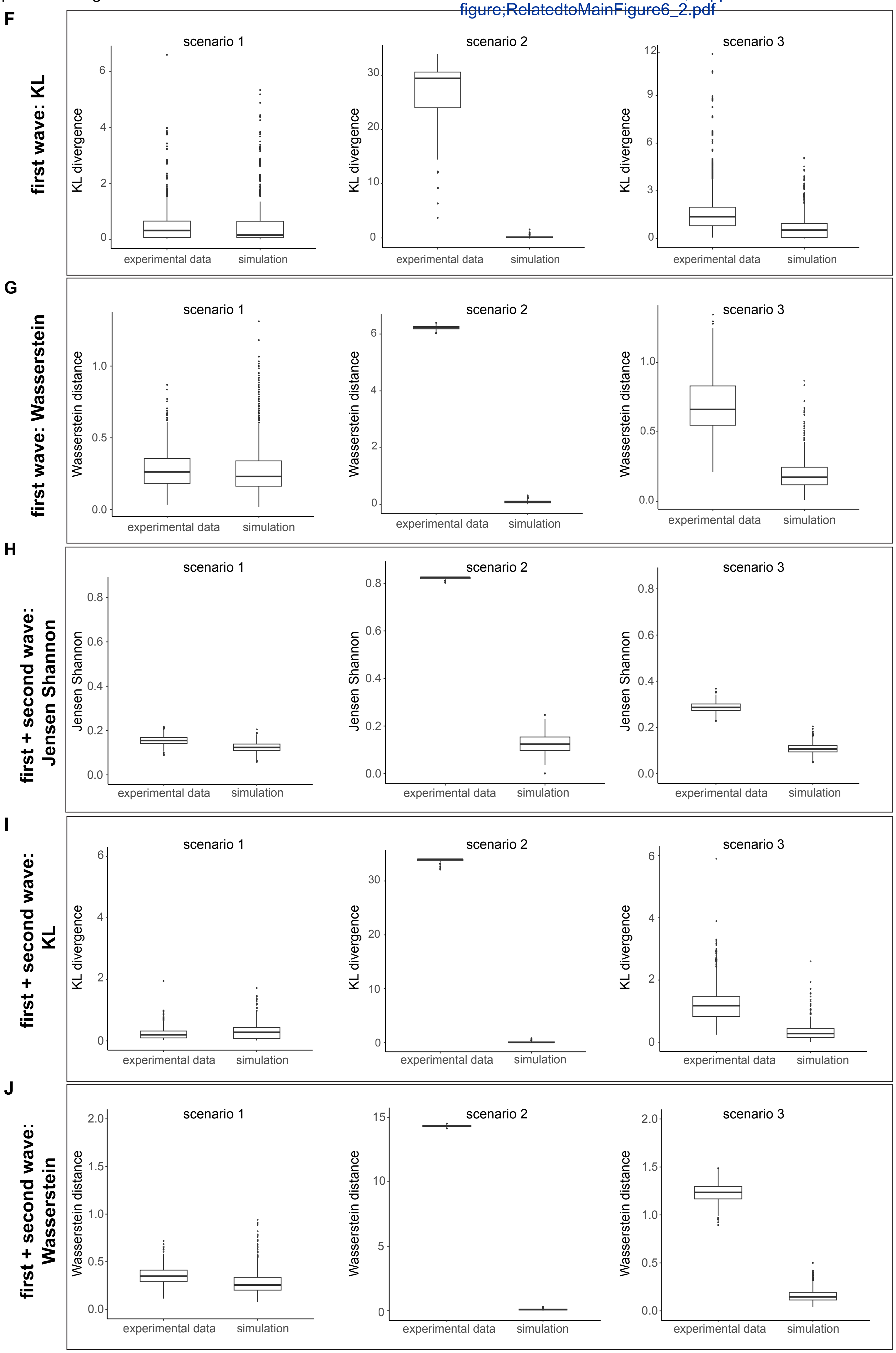

### Supplementary Figure 7

Click here to access/download;Supplemental figure;RelatedtoMainFigure7.pdf

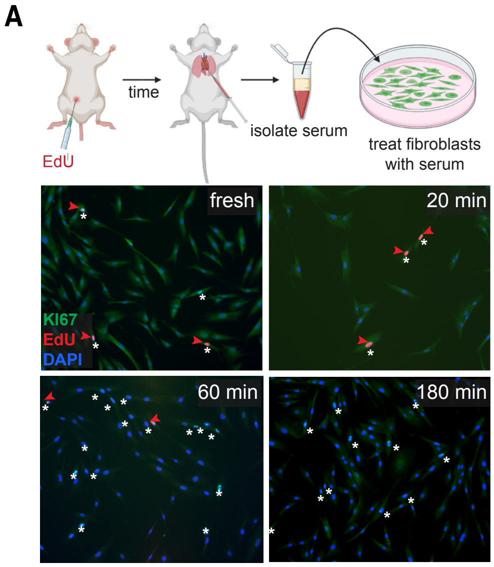


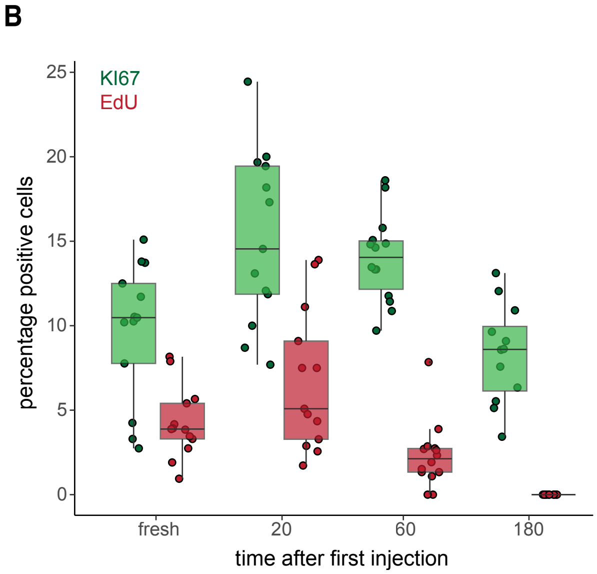

Supplemental Figure 8

Click here to access/download;Supplemental figure;RelatedtoMainFigure8_1.pdf

### Supplementary Figure 8

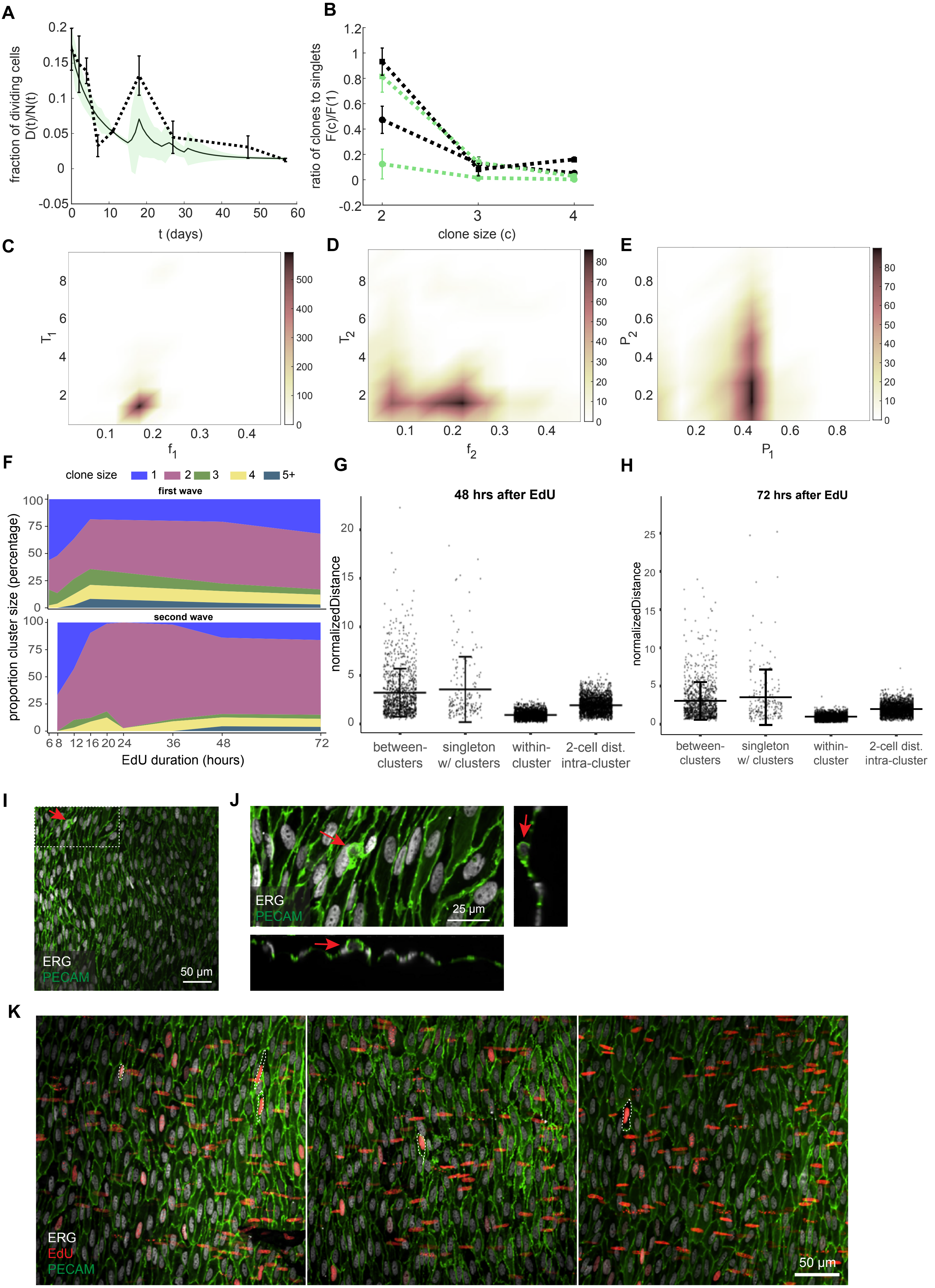

Click here to access/download;Supplemental figure;RelatedtoMainFigure8_2.pdf

### Supplementary Figure 8

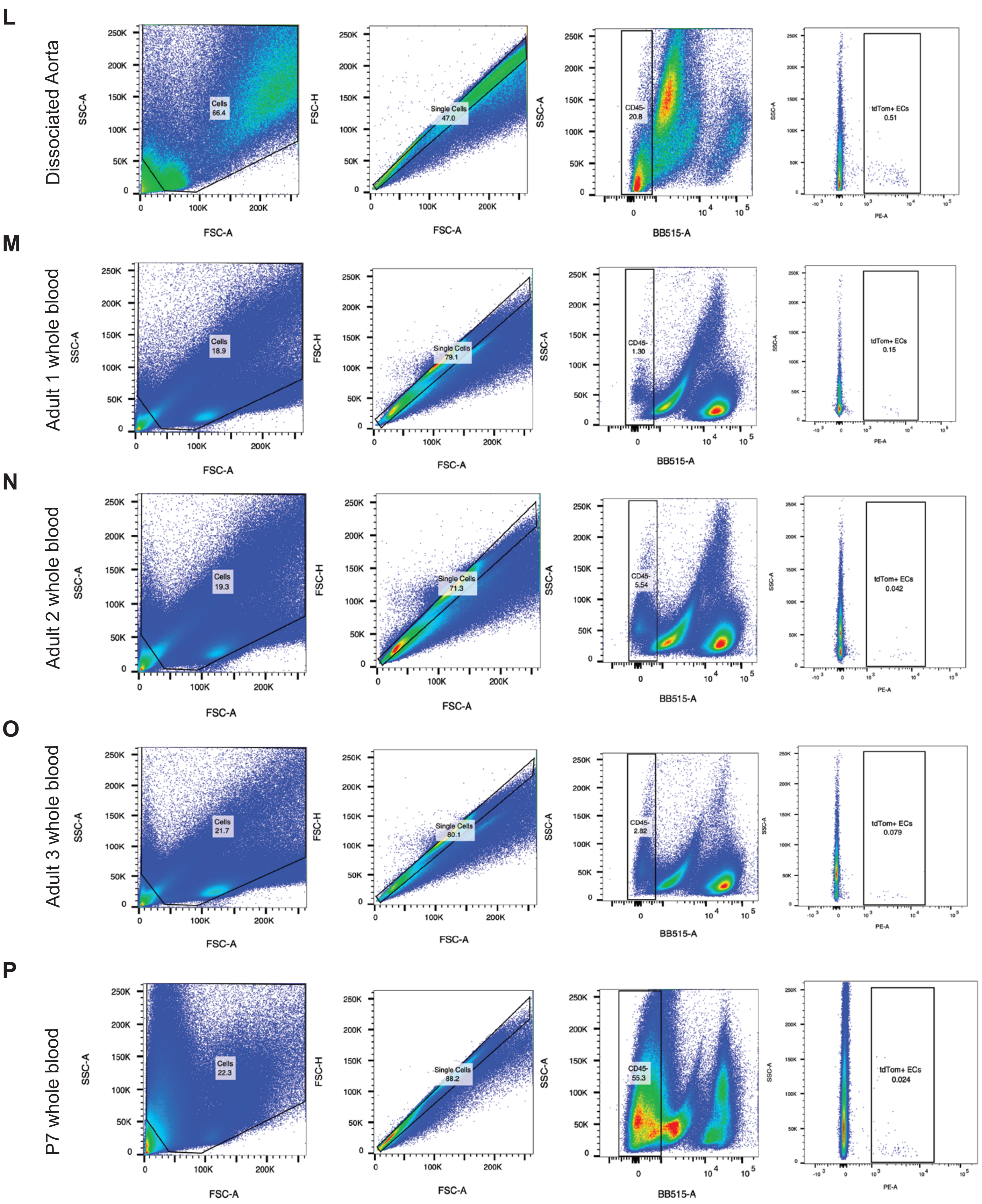

Click here to access/download;Supplemental figure;Supplementary Figure 9.pdf

**Supplementary Figure 9**

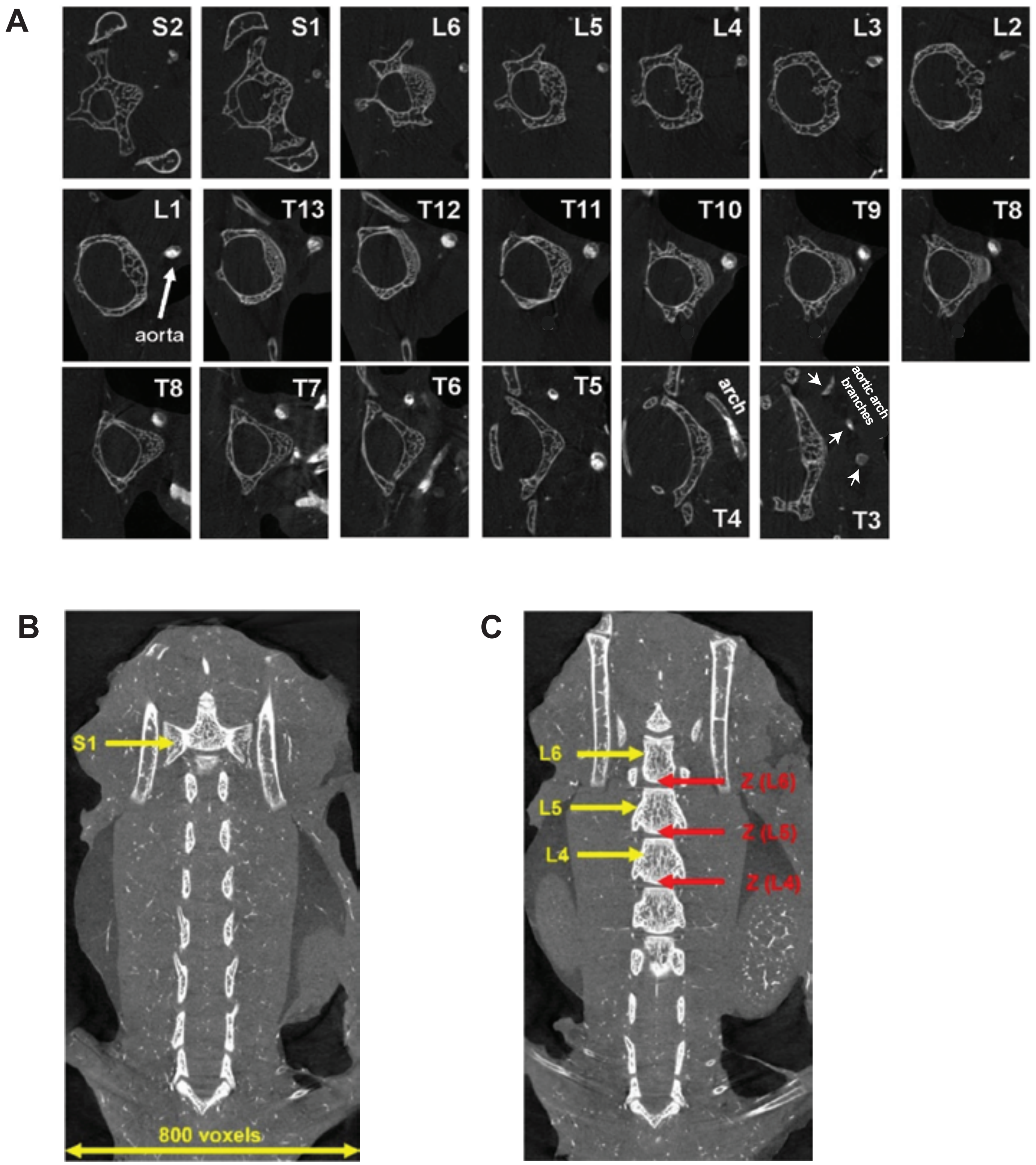
